# Spermidine/spermine N1-acetyltransferase controls tissue-specific regulatory T cell function in chronic inflammation

**DOI:** 10.1101/2024.03.25.586519

**Authors:** Teresa Neuwirth, Daniel Malzl, Katja Knapp, Panagiota Tsokkou, Lisa Kleissl, Anna Redl, Christian Freystätter, Nara Marella, Ana P. Kutschat, Elisabeth Ponweiser, Arvand Haschemi, Davide Seruggia, Jörg Menche, Erwin F. Wagner, Georg Stary

## Abstract

Regulatory T cells (T_regs_) are a critical immune component guarding against excessive inflammatory responses. During chronic inflammation, T_regs_ fail to control effector T cell responses. The causes of T_reg_ dysfunction in these diseases are poorly characterized and therapies are aimed at blocking aberrant effector responses rather than rescuing T_reg_ function. Here we utilized single-cell RNA sequencing data from patients suffering from chronic skin and colon inflammation to uncover *SAT1*, the gene encoding spermidine/spermine N1-acetyltransferase (SSAT), as a novel marker and driver of skin-specific T_reg_ dysfunction during T_H_17-mediated inflammation. T_regs_ expressing *SAT1* exhibit a tissue-specific inflammation signature and show a proinflammatory effector-like profile. In CRISPRa on healthy human skin-derived T_regs_ increased expression of *SAT1* leads to a loss of suppressive function and a switch to a T_H_17-like phenotype. This phenotype is induced by co-receptor expression on keratinocytes exposed to a T_H_17 microenvironment. Finally, the potential therapeutic impact of targeting SSAT was demonstrated in a mouse model of skin inflammation by inhibiting SSAT pharmacologically, which rescued T_reg_ number and function in the skin and systemically. Together, these data show that *SAT1* expression has severe functional consequences on T_regs_ and provides a novel target to treat chronic inflammatory skin disease.

## Introduction

Regulatory T cells (T_regs_) are a dedicated CD4^+^ T cell subset responsible for controlling excessive effector T cell responses, which is crucial in preventing unwanted and excessive immune responses^1^. Similar to other immune cell populations, T_regs_ can reside in non-lymphoid tissues in multiple different organs, where these cells display tissue-specific transcriptional programs, exhibiting T_reg_ adaptability to niche-specific regulatory requirements^2,3^. However, during chronic inflammatory diseases, such as psoriasis and inflammatory bowel disease, T_regs_ fail to inhibit the continuous immune response, which is commonly mediated by T_H_17 effector T cells^4^. This has led to the development of multiple immunotherapies targeting T_H_17-related pathways^5^. However, these therapies globally inhibit effector immune responses, increasing patients’ vulnerability to opportunistic infections^6, 7^. Hence, there would be significant advantages in reinstating the natural function of T_reg_ to regain control over effector T cells. The reasons for the failure of T_regs_ to inhibit T_H_17 function in chronic inflammation are not well understood. It has been described that T_regs_ exposed to continuous inflammatory or metabolic stimuli can become “fragile” and start producing pro-inflammatory cytokines while keeping FoxP3 expression^8, 9^. One example of metabolic control over T cell lineage commitment is polyamine metabolism, which has been shown to be important in regulating CD4^+^ helper cell stability in murine models of autoimmunity^10, 11, 12^ as well as mediating an imbalance between T_H_17 and T_regs_ in humans infected with HIV^13^. Yet, little is known about the exact mechanisms causing T_regs_ to lose their suppressive function in chronically inflamed tissue.

This study hypothesizes that T_regs_ are dysregulated in the tissue of patients affected by chronic inflammatory diseases and fail to control effector T cells and subsequent inflammation. Using the human chronic inflammatory conditions ulcerative colitis, psoriasis, and persistent cutaneous sarcoidosis as a model to compare tissue-specific T_reg_ dysfunction we sought to investigate why T_regs_ fail to control effector T cells in a T_H_17 environment. The data shown here demonstrate that spermidine/spermine N1-acetyltransferase (SSAT), an enzyme of the polyamine metabolic pathway, is responsible for T_reg_ suppressive function in the skin and controls the switch between functionally intact T_reg_ and a fragile, T_H_17-like T_reg_ identity. These data provide mechanistic insights in the dysregulation of human T_regs_ in an inflammatory environment in the skin by losing their canonical regulatory function and suggest a novel approach towards T_reg_-targeted therapies for chronic inflammation.

## Results

### T_regs_ are reduced in chronic inflammation exhibiting a tissue-specific inflammation signature

First, we assessed the similarity of T_regs_ on a transcriptional level in different organ-confined, T_H_17-mediated chronic inflammatory diseases. To investigate tissue-specific signatures of T_regs_, we compared single-cell RNA sequencing (scRNAseq) datasets of chronic inflammatory diseases affecting the skin or the colon. Using publicly available datasets for psoriasis, persistent cutaneous sarcoidosis and ulcerative colitis^14, 15, 16^, we employed a common processing workflow for all datasets to mitigate any processing-related errors introduced by different data handling and to ensure that sampling– or patient-specific variation was removed prior to cell type identification (Extended Data Fig. 1A). Starting from the raw sequence data, the raw count data was projected into a common embedding space using scVI^17^. In total, we integrated data of 3 psoriasis^15^ and 14 sarcoidosis patients^14^ with 5 healthy volunteers^15, 18^, clustered the cells and extracted T cell clusters on the basis of CD3D expression (Extended Data Fig. 1B). In parallel, a similar workflow was applied to a previously published dataset of ulcerative colitis (UC)^16^ consisting of 7 patients and 8 healthy volunteer samples of intestinal tissue (Extended Data Fig 1B). In both skin and colon, we identified clusters of T_regs_ based on differential expression of *FOXP3, IL2RA,* and *IKZF2* (Fig. 1A and B, Extended Data Fig. 1C). To evaluate a potential reduction of T_regs_, we then assessed the abundance of T_regs_ in the scRNA-seq data. Using differential cell abundance (DCA) testing based on k-nearest neighbor (kNN) graphs^19^, T_reg_ counts were reduced in chronic cutaneous inflammation compared to healthy tissue (Fig. 1A), which was not observed in the UC dataset (Fig 1B). To validate these findings, T_regs_ were stained in skin and colon sections from patients and healthy controls. This showed a mild influx of T cells in samples from sarcoidosis and psoriasis patients and comparable T cell numbers in UC patients and healthy colon tissue (Fig. 1C). However, a significant reduction of percentage of T_regs_ compared to healthy individuals was observed in patient samples in both skin and colon (Fig. 1D and E). While this confirmed the observation for a lower abundance of T_regs_ at RNA level from the skin, no significant changes in the prevalence of T_regs_ in the UC scRNAseq dataset were observed. This discrepancy in colon samples could have multiple reasons, including differences in tissue processing between IF and scRNAseq, changes depending on the exact sampling site or diverse disease profiles as samples from different patients were used for immunofluorescence staining and scRNAseq.

**Figure 1:**
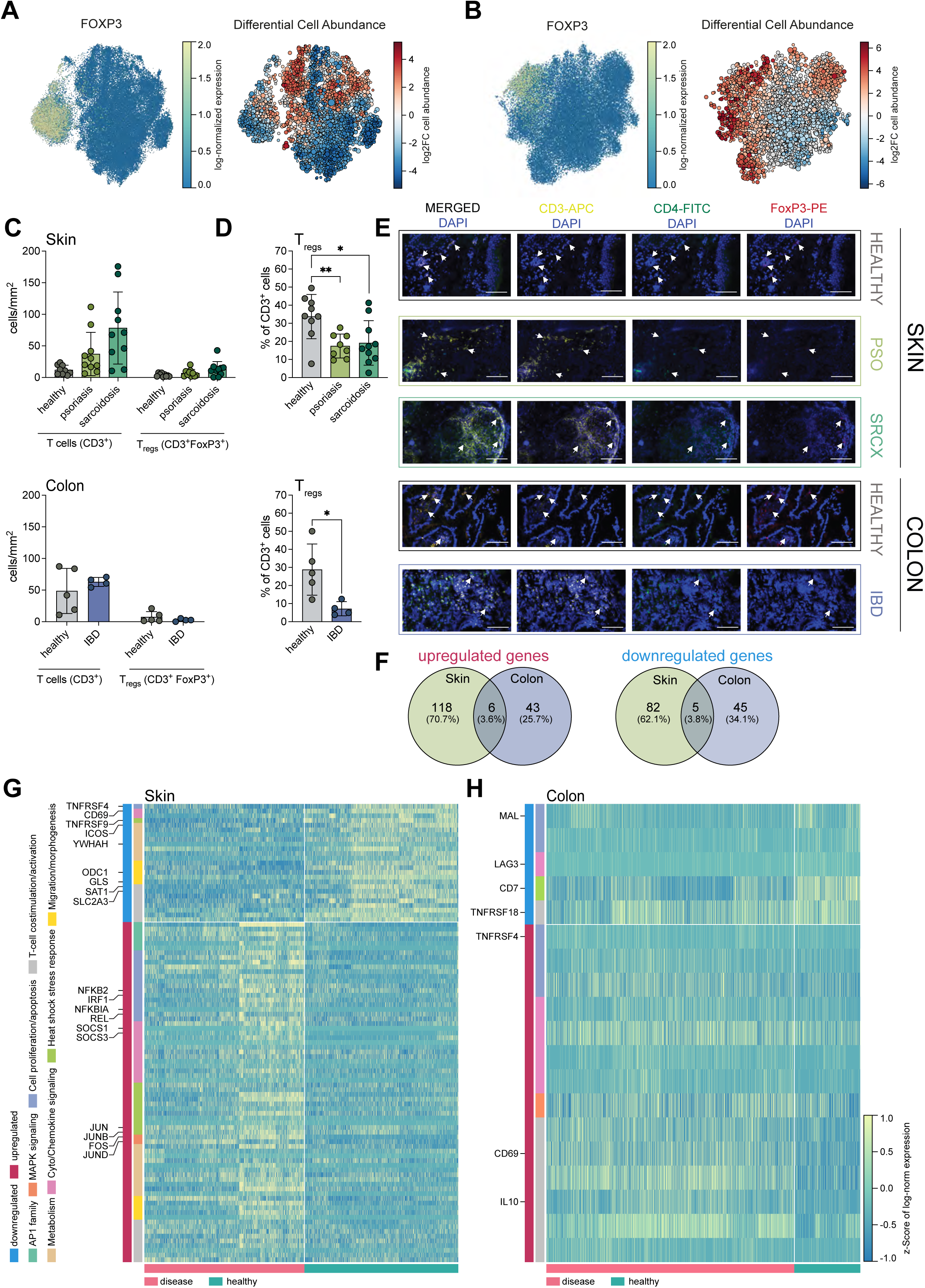
T_regs_ are reduced in chronic inflammation and show a tissue-specific inflammation signature. **A)** and **B)** UMAP projection of FoxP3 expression and differential abundance analysis of (A) skin T cells including healthy (n=5), psoriasis (n=3), and sarcoidosis (n=14) samples and (B) healthy colon (n=8) and ulcerative colitis samples (n=7). **C)** and **D)** Quantification of immunofluorescence staining for T cells (CD3^+^) and T_regs_ (CD3^+^FoxP3^+^) in skin and colon samples as (C) absolute numbers/mm^2^ and (D) percentage of T cells. Each data point represents one patient or healthy donor. Mean ± SD. **E)** Representative images of T_reg_ staining in patient and healthy tissue. Scale bar = 100μm. **F)** Overlap of differentially expressed genes between T_regs_ in skin and colon inflammation. **G)** and **H)** Heatmap of a selection of differentially expressed genes between T_regs_ in (G) healthy and inflamed skin and (H) healthy and inflamed colon. Two-way ANOVA with Holm-Sidak multiple-testing correction, *p<0.05, **p<0.01.

To further investigate T_reg_ signatures in inflamed tissue, we performed differential gene expression analysis (DEA) to identify differentially expressed genes (DEGs) in healthy versus diseased samples. Differences in gene expression were more pronounced in skin compared to colon T_regs_ (211 DEGs vs. 101 DEGs), indicating that T_regs_ in the skin are more affected by or involved in the pathogenesis of T_H_17-inflammation than in the colon. Overall, there was a low overlap in DEGs between the skin and the colon under inflammatory conditions (6 genes shared upregulated: *LTB, DUSP2, MAGEH, TNFRSF4, RELB, PDE4D*; 6 shared downregulated: *TXNIP, MT-ND3, LGALS1, RPL36A, RPL31*) (Fig. 1F, G and H), emphasizing that T_reg_ signatures are specific for tissues rather than shared in chronic inflammation. T_regs_ derived from inflamed skin showed, among others, an upregulation of AP1 family members (*FOS, JUN, JUNB, JUND*), regulation of cytokine signaling (*SOCS3, SOCS1, NFKBIA, IRF1, NFKB2, REL*), T cell co-stimulation and activation (*ICOS, CD69, TNFRSF9, TNFRSF4, YWHAH*), and metabolism (*SAT1, ODC1, SLC2A3, GLS*) (Fig. 1G). Overall, the difference in T_regs_ from inflamed versus healthy colon were less pronounced with an upregulation of T cell co-stimulatory and activation markers (*TNFRSF4, TNFRSF18, CD7, LAG3, MAL*), and a downregulation of *CD69* and *IL10* (Fig. 1H). These results emphasize that, while psoriasis, sarcoidosis, and ulcerative colitis are considered T_H_17-driven diseases, T_regs_ are affected differently in various organs, indicating that failure to regulate pathogenic T cell responses in chronic inflammation has underlying mechanisms specific for the skin and colon.

### Expression of SSAT defines a T_reg_ subset specific for cutaneous inflammation

While T_regs_ are depleted in the skin and colon upon chronic inflammation, DCA analysis showed an enrichment of a small T_reg_ subpopulation in inflamed skin. Therefore, we performed gene ranking of enriched neighborhoods within this T_reg_ cluster. Looking at the top 5 ranked genes, *SAT1* caught our interest (Extended Data Fig. 2A). This gene was upregulated in T_regs_ in skin under inflammatory conditions and mapped particularly well to the enriched T_reg_ cluster, dividing T_regs_ into 2 populations: SAT1^high^ and SAT1^low^ (Fig. 2A-D). *SAT1* encodes SSAT, which is a rate limiting enzyme of the polyamine catabolic pathway. Polyamines have recently been shown to affect CD4^+^ effector T cell function and lineage stability^10, 11, 12^, although mechanistic studies have so far been limited to mouse models for autoimmunity. Therefore, this gene was chosen for investigating its influence on human T_reg_ function. *SAT1* expression was highest in skin T_regs_ under chronic inflammation (Extended Data Fig. 2B) and, within the T cell compartment, its high expression in disease was specific to T_regs_ (Fig. 2E and Extended Data Fig 2C).

**Figure 2:**
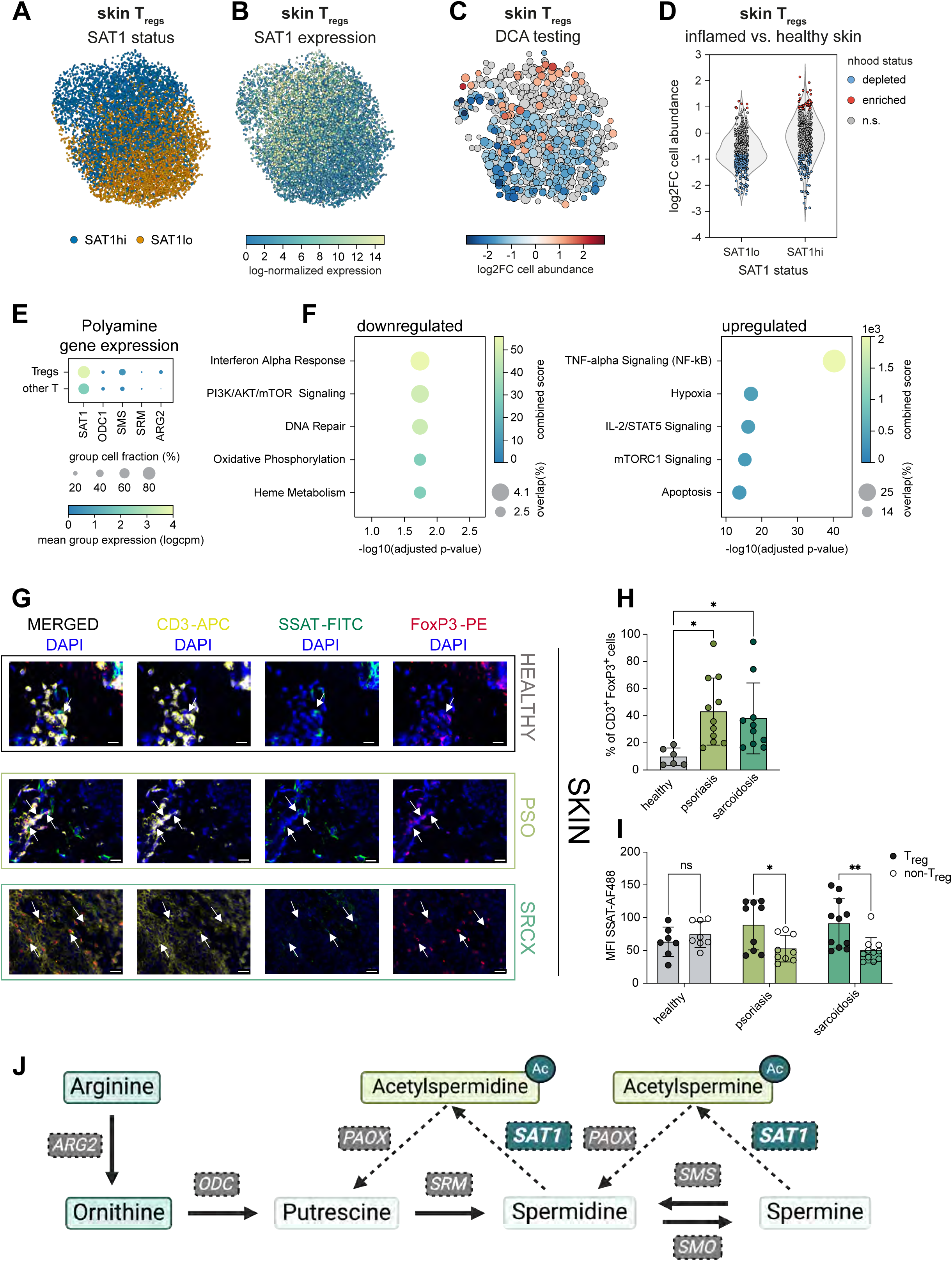
S*A*T1 expression defines an inflammation-specific T_reg_ population in the skin. **A)** UMAP projection of skin T_regs_ showing distribution of SAT1^high^ and SAT1^low^ cells. **B)** UMAP projection of skin T_regs_ showing *SAT1* expression. **C)** Differential abundance testing of skin T_regs_ in inflamed vs. healthy skin. **D)** Violin plot of log2-fold changes of neighborhood cell counts in differential abundance testing of SAT1^high^ and SAT1^low^ T_regs_ in the skin. **E)** Dotplot of expression of genes involved in polyamine metabolism in skin T_regs_ and non-T_reg_ T cells. **F)** Gene set enrichment analysis of differentially expressed genes between SAT1^high^ vs. SAT1^low^ skin T_regs_. Shown are the top 5 enriched MSigDB Hallmark gene sets for up– and downregulated genes. **G)** Representative images of SSAT staining in T cells in the skin. Scale bar = 20μm. **H)** Quantification of immunofluorescence staining of SSAT^+^ T_regs_ (CD3^+^ FoxP3^+^) in the skin. **I)** Mean fluorescence intensity of SSAT-AF488 immunofluorescence staining in T_regs_ (CD3^+^FoxP3^+^) and non-T_reg_ T cells (CD3^+^) in the skin. **J)** Graphic of the polyamine metabolic pathway. Created using BioRender. **H)** and **I)** Each dot represents one patient or healthy donor. Mean ± SD. Two-way ANOVA with Holm-Sidak multiple-testing correction, ns p>0.05, *p<0.05.

To directly compare SAT1^high^ and SAT1^low^ T_reg_ populations of the skin, we performed gene set enrichment analysis (GSEA) of genes differentially expressed in SAT1^high^ versus SAT1^low^ T_regs_ in chronic cutaneous inflammation (Fig. 2F). Hallmark terms enriched in SAT1^high^ T_regs_ include hypoxia, as well as TNF-α signaling, a known regulator of SSAT transcription^20^. Terms enriched in downregulated genes included oxidative phosphorylation and mTOR signaling via PI3K. This indicates an imbalance in nutrient utilization (mTOR signaling) and metabolic fitness of SAT1^high^ T_regs_. Using immunofluorescence staining of skin cryosections, expression of SSAT on protein level in T cells and T_regs_ in psoriasis and sarcoidosis skin was validated (Fig. 2G), confirming that SSAT^+^ T_regs_ are more abundant in diseased skin (Fig. 2H). Furthermore, T_regs_ in psoriasis and sarcoidosis had a significantly higher mean fluorescence intensity (MFI) in SSAT^+^ compared to other T cells, which was not the case in healthy skin (Fig. 2I). Of note, there was no enrichment of SSAT^+^ T_regs_ in colon biopsies from IBD patients (Extended Data Fig. 2E-G). Overall, immunofluorescence staining of SSAT in T_regs_ confirmed the observations from the scRNAseq data at protein level with SAT1^high^ T_regs_ occurring in the skin upon chronic inflammation.

The main cellular function of SSAT is to acetylate lower polyamines, which can then be either fed back into the catabolic pathway via oxidation or be exported from cells in their acetylated form (Fig. 2J). Therefore, we hypothesized that T_regs_ isolated from chronic inflammatory skin diseases would most likely have a decrease in intracellular polyamine levels. While SAT1^high^ T_regs_ in the blood of psoriasis and sarcoidosis patients were less abundant as opposed to the skin, there was a T_reg_ population expressing *SAT1* (Extended Data Fig. 3A-E). To avoid extensive digestion protocols of skin and reach desired cell numbers for LC-MS metabolomics, we isolated blood-derived T_regs_ from psoriasis and sarcoidosis patients using magnetic beads, yielding pure T_reg_ populations (Extended Data Fig. 3F). Spermidine and spermine were significantly decreased (approx. 10%) in T_regs_ from psoriasis and sarcoidosis patients compared to healthy control subjects, further emphasizing that SAT1^high^ T_regs_ have a change in polyamine metabolism, which could affect their function. (Extended Data Fig. 3G, Supplementary Table 1).

To investigate whether upregulation of *SAT1* is specific to diseases driven by T_H_17– inflammation, we then analyzed data from atopic dermatitis^15^ (AD) (4 patients, 8 healthy volunteers), a well-described T_H_2-drived disease (Extended Data Fig. 4A-E). In these patients, there was a slight depletion of T_regs_ based on differential abundance testing (Extended Data Fig. 4F). Like in T_H_17 inflammation, there was a significant increase in *SAT1* expression in T_regs_ (but not other T cells) from AD patients (Extended Data Fig. 4G and H). While AD is traditionally associated with a T_H_2 response, some studies suggest that there may also be a T_H_17 component to the disease^21, 22, 23^. We used T_H_17– and T_H_2-associated genes (see Methods) to calculate a T_H_17 and T_H_2 score, respectively. Indeed, there was a pronounced signature of T_H_17 inflammation also in AD patients (Extended Data Fig. 4I and J), explaining why this signature is also observed in AD. These results show that high expression of the polyamine acetyltransferase SSAT is a hallmark of dysfunctional T_regs_ in chronically inflamed skin and associated with a T_H_17 signature.

### *SAT1* expression impairs the phenotype and function of T_regs_

Given the association of SAT1^high^ T_regs_ with a dysregulated phenotype in chronic inflammation, we then sought to investigate the functional consequences of increased *SAT1* expression. T_regs_ were isolated from healthy skin donors, expressed *SAT1* with a CRISPRa-mediated system and assessed their phenotype and function (Fig. 3A). Bulk T cells were first expanded over two weeks using a skin explant and T_regs_ were subsequently isolated with magnetic beads based on CD25^hi^ and CD127^lo^ expression in CD4^+^ T cells (Extended Data Fig. 3F and Extended Data Fig. 5A). The CRISPRa-mediated expression system achieved an approx. 10-fold increase in expression of *SAT1* in skin-derived T_regs_ (Extended Data Fig. 5B and 5C). Using surface markers differentially expressed between SAT1^high^ and SAT1^low^ T_regs_ recapitulated the phenotype of SAT1^high^ T_regs_ from the scRNA-seq data (Supplementary Table 2; referred to as aSAT1 T_regs_ from here on). Compared to T_regs_ transduced with a non-targeting (NT) control guide, aT_regs_ showed a significant upregulation of the skin-residency and activation markers CXCR6 and CD69 as well as the co-inhibitory molecule PD-1 by aSAT1 T_regs_ (Fig. 3B-D). All these molecules are important for T_reg_ function and high PD-1 expression has previously been associated with a dysfunctional T_reg_ phenotype^24^, which led us to investigate cytokine production by these cells. aSAT1 T_regs_ produce significantly more IL-17A and IL-2 (Fig. 3E-G) than cells transduced with a non-targeting guide, indicating that high expression of *SAT1* in T_regs_ shifts their phenotype towards T_H_17 effector T cells. In line with these observations, the T_H_17-lineage transcription factor ROR-γt was expressed higher in aSAT1 T_regs_, while FoxP3 expression remained unchanged (Fig. 3H-J). CXCR6, CD69 and cytokines (IL-17A, IL-2, IL-10) remained unchanged in aSAT1 CD4^+^ T_eff_ cells, indicating that upregulation of these molecules in response to *SAT1* expression is T_reg_ specific (Extended Data Fig. 5D). As this conversion from T_reg_ to a T_H_17-like phenotype was observed upon *SAT1* expression *in vitro*, we used a gene signature usually expressed by effector CD4^+^ T cells to investigate whether SAT1^high^ T_regs_ are more like “fragile T_regs_” (i.e. T_regs_ that lose their regulatory phenotype but keep FoxP3 expression, see Methods) and applied it to SAT1^high^ and SAT1l^ow^ T_regs_ from the scRNAseq data. This score confirmed that SAT1^high^ T_regs_ have an increased “fragile T_reg_” signature compared to SAT1^low^ T_regs_ (Fig. 3K), similar to what was observed upon *SAT1* expression *in vitro* via CRISPRa.

**Figure 3:**
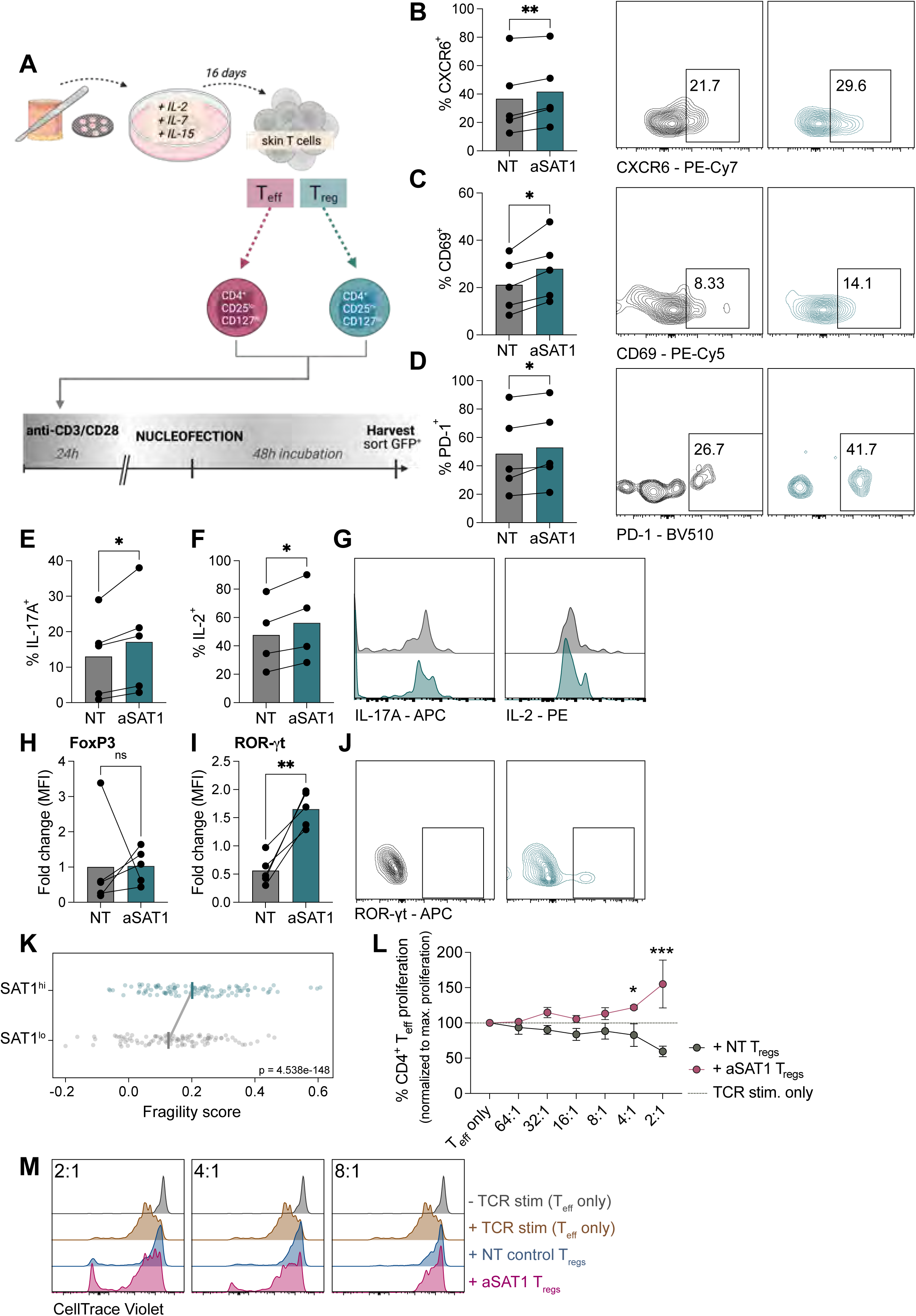
SAT1high T_regs_ exhibit an effector phenotype and fail to inhibit T cell proliferation. **A)** Experimental design using skin-derived T cells for CRISPRa-mediated expression of *SAT1*. **B)-F)** Quantification of flow cytometry data showing percentage of (B) CXCR6^+,^ (C) CD69^+^, (**D**) PD-1^+^, and flow cytometry density plot of representative data (right) and (E) IL-17A^+^, (**F**) IL-2^+^ cells in GFP^+^ T_regs_ in non-targeting vs. SAT1 transduced cells. **G)** Representative histograms of IL-17A and IL-2 expression in non-targeting vs. SAT1 transduced T_regs_. **H)** and **I)** Quantification of flow cytometry data showing fold change in mean fluorescence intensity (MFI) of (H) FoxP3 and (I) ROR-γt in non-targeting vs. SAT1 transduced T_regs_. Fold change was calculated relative to non-targeting control cells. **J)** Representative flow cytometry density plot of ROR-γt expression in non-targeting vs. SAT1 transduced T_regs_. **K)** Fragility score of SAT1^high^ and SAT1^low^ T_regs_ computed as the ratio of average expression of target genes over a random background (see methods). Lines are median score of all cells in category. Dots are a random sample of 100 cells for each category. **L)** Quantification of flow cytometry based proliferation readout of CellTrace Violet labelled CD4^+^ T_eff_ cells co-cultured with non-targeting or SAT1 transduced T_regs_ at varying ratio for 5 days. Percentage proliferation shown as normalized proliferation to respective maximum proliferation with TCR stimulation only. **M)** Representative histograms showing proliferation based on CellTrace Violet of CD4^+^ T_eff_ co-cultured with non-targeting or SAT1 transduced T_regs_ at varying ratio for 5 days. **B)-H)** n=5 healthy skin donors, Two-tailed paired t-test, ns p>0.05, *p<0.05, **p<0.01. **K)** p-Value based on two-tailed t-test. **L)** n=4 healthy skin donors, Two-way ANOVA with Holm-Sidak multiple-testing correction, ns p>0.05, *p<0.05, **p<0.01, ***p<0.001.

A phenotypical switch of stable, suppressive T_regs_ to fragile T_regs_ could explain a loss of T_reg_ function in chronic skin inflammation. As such, we next investigated the suppressive capacity of aSAT1 T_regs_ in order to test the possibility that increased expression of *SAT1* could explain the lack of immunosuppression seen in patients. Therefore, we co-cultured CD4^+^ effector T cells (CD25^low^ CD127^high^) with aSAT1 or NT control T_regs_ at increasing ratios. After 5 days of co-culture and TCR stimulation, aSAT1 T_regs_ failed to inhibit CD4^+^ T_eff_ cell proliferation and actually drove their expansion (Fig. 3L and M). These results indicate that a high expression of *SAT1* has severe functional consequences in T_regs_, which could contribute to excessive T_eff_ activation described in chronic skin inflammation.

### SAT1^high^ T_regs_ are induced by keratinocytes in a T_H_17 environment

Given that expression of *SAT1* impairs cell function and phenotype of skin-derived T_regs_, we next investigated the role of the tissue microenvironment for the induction of SAT1^high^ T_regs_. The function of T cells in tissue is highly dependent on environmental cues and interactions with different cell types. Therefore, we used CellChat to assess the scRNAseq dataset of psoriasis and sarcoidosis skin for signals received by SAT1^high^ and SAT1^low^ T_regs_ from different cell populations in the skin. T_regs_ had incoming signals related to adhesion and migration (NECTIN, GALECTIN) and SAT1^high^ T_regs_ showed especially strong enrichments in T cell co-stimulation (CD137, CD86, CD80, OX40) as well as cytokine and chemokine signaling (CCL, TNF; Fig. 4A).

**Figure 4:**
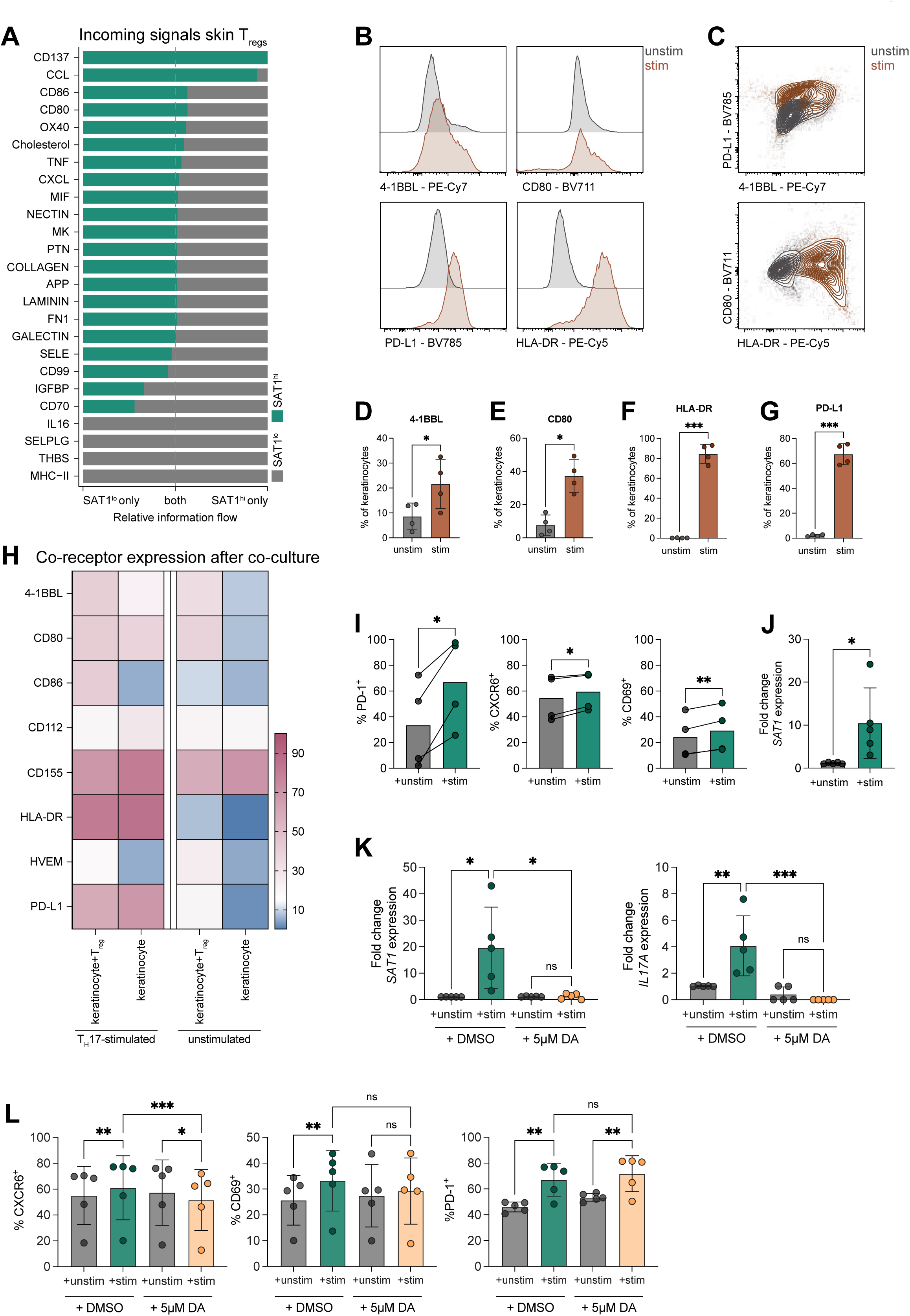
Keratinocytes exposed to a T_H_17 inflammatory environment drive SAT1^high^ phenotype in T_regs_. **A)** Interaction analysis of fraction-based incoming signals in SAT1^high^ and SAT1^low^ T_regs_ in chronic skin inflammation. **B)** Representative histograms of expression of 4-1BBL, CD80, PD-L1, and HLA-DR in keratinocytes after overnight stimulation with T_H_17 cytokines (IL-17A, IFN-γ, and IL-1β) and unstimulated controls. **C)** Representative flow cytometry density plot of co-expression of co-receptors by keratinocytes after overnight stimulation with T_H_17 cytokines and unstimulated controls. **D)-G)** Quantification of flow cytometry data showing percentage of (D) 4-1BBL^+^, (E) CD80^+^, (F) HLA-DR^+^, (G) PD-L^+^ in keratinocytes either stimulated with T_H_17 cytokines or media-only. **H)** Heatmap of flow cytometry data showing expression of co-receptors by keratinocytes after T_H_17 stimulation and co-culture with healthy blood-derived T_regs_. Scale represents percent of expressing cells. **I)** Quantification of flow cytometry data showing percentage of PD-1^+^, CXCR6^+^, and CD69^+^ blood-derived T_regs_ after co-culture with T_H_17 stimulated or unstimulated keratinocytes for 5 days. **J)** Fold change of *SAT1* expression in healthy, blood-derived T_regs_ co-cultured with T_H_17 or unstimulated keratinocytes for 5 days measured by qPCR. Fold change calculated relative to unstimulated control. **K)** Fold change of *SAT1* (left) and *IL17A* (right) expression in healthy, blood-derived T_regs_ co-cultured with T_H_17 or unstimulated keratinocytes for 5 days with and without diminazene aceturate (SSAT inhibitor) measured by qPCR. Fold change calculated relative to each unstimulated control. **L)** Quantification of flow cytometry data showing percentage of PD-1^+^, CXCR6^+^, and CD69^+^ blood-derived T_regs_ after co-culture with T_H_17 stimulated or unstimulated keratinocytes for 5 days with and without diminazene aceturate. **D)-G)** Two-tailed unpaired t-test, *p<0.05, **p<0.01, ***p<0.001. Mean ± SD. **I)-J)** Two-tailed paired t-test, *p<0.05, **p<0.01. **K)-L)** Two-way ANOVA with Tukey correction for multiple testing, *p<0.05, **p<0.01, ***p<0.001. Mean ± SD.

Upregulation of residency markers, such as CXCR6 has previously been described on T cells with homing potential to non-lymphoid tissue and skin residency development^25,26^. Thus, we we then investigated if incoming T_regs_ from the circulation could induce *SAT1* expression by the local inflammatory microenvironment and therefore fail to break the inflammatory loop. Keratinocytes are known to be of importance in driving and inducing T_H_17-inflammation in the skin^27^ and are also able to express T cell co-stimulatory molecules^28, 29, 30, 31^. Hence, we hypothesized that keratinocytes exposed to a T_H_17-environment could drive a pathogenic SAT1^high^ T_reg_ phenotype. To test this hypothesis, keratinocytes were stimulated with a T_H_17 cytokine cocktail and assessed induction of co-receptors observed in the interaction data. Indeed, after stimulation keratinocytes significantly increased expression of 4-1BBL (CD137L), CD80, HLA-DR, and PD-L1 and these cells tended to co-express multiple of these receptors (Fig. 4B-G). We then isolated T_regs_ from the blood of healthy donors and co-cultured these cells with “inflammation-primed” keratinocytes. Co-culture with T cells slightly upregulated expression of multiple co-receptors on keratinocytes even without stimulation, but overall upregulation was more pronounced with T_H_17-cytokine stimulation (Fig. 4H, and Extended Data Fig. 6C). However, co-culture with unstimulated keratinocytes did not significantly alter SAT1^high^ T_reg_ markers in T_regs_ or T_eff_ compared to culture without keratinocytes (Extended Data Fig. 6A and B). After 5 days of co-culture, T_regs_ exposed to T_H_17-cytokine-primed keratinocytes expressed higher amounts of PD-1, CXCR6, and CD69 as measured by flow cytometry (Fig. 4I). Furthermore, these cells also upregulated RNA expression of *SAT1*, showing that keratinocytes exposed to T_H_17 stimuli can induce a SAT1^high^ phenotype in healthy T_regs_ (Fig. 4J). Upregulation of PD-1 was also observed in a co-culture with T_eff_, however CXCR6 was downregulated, and no significant change was observed in CD69 expression on T_eff_ (Extended Data Fig. 6D). Induction of *SAT1* expression was only seen in T_regs_ and no significant change in gene expression was seen in co-cultured T_eff_ (Extended Data Fig. 6E), suggesting that co-receptor expression in a T_H_17-inflammatory environment specifically induced the expression of *SAT1* and a subsequent phenotype switch in T_regs_.

Given that upregulation of *SAT1* in T_regs_ was also observed in AD patients, we next investigated if keratinocytes exposed to a “clean” T_H_2 environment (i.e., only IL-4 with no T_H_17 background) could also upregulate co-receptors or if this is a T_H_17-specific effect. Consistently, there was no induction of co-receptor expression in keratinocytes after stimulation with IL-4 (Extended Data Fig. 7A) or induction of a SAT1^high^ phenotype in T cells co-cultured with T_H_2-stimulated keratinocytes (Extended Data Fig. 7B), further supporting the hypothesis that this phenotype is driven by the microenvironment of T_H_17-associated cutaneous inflammation. Taken together, these data show that *SAT1* gene expression and the resulting “fragile” phenotype of Tregs can be induced by keratinocytes exposed to a T_H_17 environment which is, at least partly, driven by co-receptor stimulation.

### Inhibition of SSAT inhibits T_H_17-like phenotype in T_regs_

To test if the induction of *SAT1* is directly responsible for the induction of the observed phenotype, SSAT was blocked using diminazene aceturate (DA), a known competitive inhibitor of SSAT ^32^. DA did not significantly alter cell viability or co-receptor expression by keratinocytes at low concentrations (1-20μM) (Extended Data Fig. 8). Therefore, a concentration of 5μM was chosen for experiments to exclude the possibility that the observed effect is mediated by keratinocytes rather than directly on T_regs_. Blood-derived T_regs_ were co-cultured with keratinocytes as described previously, and either DMSO or DA was added to the co-cultures. Inhibiting SSAT using DA led to a reduction of *SAT1* gene expression as well as blocking of the induction of *IL17A* expression (Fig. 4K). Furthermore, CXCR6 and CD69 expression were inhibited by DA, but there was no effect on PD-1 expression. This indicates that PD-1 is most likely regulated independently of SSAT (Fig. 4L), while CXCR6 and CD69 seem to be directly affected by the expression of *SAT1* and activity of SSAT. This data demonstrates that SSAT induces a pathogenic phenotype in T_regs_ upregulating tissue-resident T cell markers upon exposure to a T_H_17-polarized microenvironment and that blocking this enzyme rescued this phenotype.

### SSAT inhibition reverses T_reg_ reduction and function in a mouse model for psoriasis

Given that expression of *SAT1* and inhibition of its product SSAT controls T_reg_ function *in vitro*, we next investigated whether targeting this enzyme has potential therapeutic impact in an *in vivo* model of chronic skin inflammation. The genetically engineered mouse model of inducible loss-of function of c-Jun and JunB transcription factors in the epidermis (DKO*^K15^)^33^ is an established psoriasis-like disease model which closely recapitulates our *in vitro* conditions. Using the DKO*^K15^ mice, we fist validated the reduction of T_regs_ and a high SSAT expression in chronically inflamed mouse skin. Immunofluorescence staining showed a significant decrease of T_reg_ numbers in the skin in DKO*^K15^ mice compared to their wild-type (WT) littermates (Fig. 5A and Extended Data Fig. 9B). As there was no commercially available antibody cross-reactive with mouse SSAT, we used a biochemical approach to stain for the activity of SSAT in the tissue. This was based on enzyme histochemical approaches described by Miller et al.^34^ and adapted for SSAT activity. SSAT enzyme activity was stained and measured by detecting released coenzyme A (coA) from acetyl-CoA in an activity assay using SSAT-specific substrates spermine and spermidine (Extended Data Fig. 9A). Enzyme activity was measured and reported as relative MFI over SSAT inhibitor-treated slides (where the inhibitor is DA) to ensure signal specificity. These stainings showed a global increase in SSAT activity in the skin of DKO*^K15^ mice compared to WT control (Fig. 5B and C). As DKO*^K^^15^ mice had reduced levels of T_regs_ and an increase of SSAT activity, this mouse model for psoriasiform skin inflammation sufficiently mimicked the human data and these mice were chosen for further experiments.

**Figure 5:**
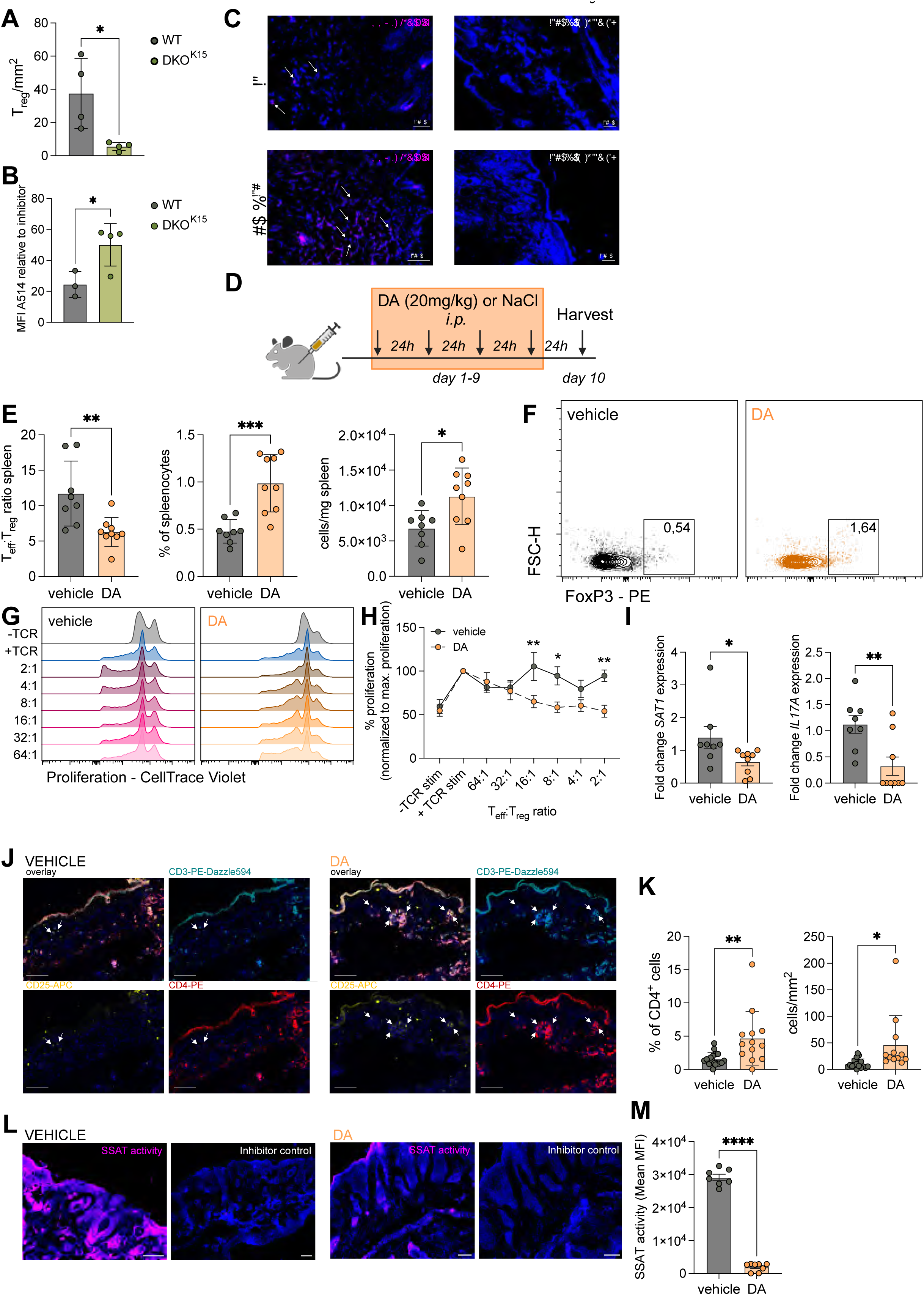
Inhibition of SSAT in a mouse model of psoriasis rescues T_reg_ number and function. **A)** Quantification of immunofluorescence staining for T_regs_ (CD3^+^CD4^+^CD25^++^) in skin of DKO*^K15^ mice as absolute numbers/mm^2^. **B)** Quantification of mean fluorescence intensity of SSAT activity in the skin of DKO*^K15^ mice. **C)** Representative images of SSAT activity staining in mouse skin. Inhibitor control = addition of diminazene aceturate to measure specific activity. Scale bar = 50μm. **D)** Experimental design for treatment of DKO*^K15^ mice with the SSAT inhibitor diminazene aceturate. **E)** CD4^+^ T_eff_ to T_reg_ ratio, percentage of T_regs_, and absolute numbers of T_regs_ in the spleen of mice after 10 days with and without treatment. **F)** Representative flow cytometry data showing expression of FoxP3 in spleen-derived T cells (gated on CD3^+^CD4^+^ cells). **G)** Representative histogram showing proliferation based on CellTrace Violet of CD4^+^ T_eff_ co-cultured with treated or untreated spleen-derived T_regs_ at varying ratio for 5 days. **H)** Quantification of flow cytometry based proliferation readout of CellTrace Violet labelled CD4^+^ T^eff^ cells co-cultured with treated or untreated spleen-derived T_regs_ at varying ratio for 5 days. Percentage proliferation shown as normalized proliferation to respective maximum proliferation with TCR stimulation only. **I)** Fold change of *SAT1* (left) and *IL17A* (right) expression in treated and untreated spleen-derived T_regs_ measured by qPCR. Fold change calculated relative to untreated control. **J)** Representative images of immunofluorescence staining for T_regs_ in mouse skin with and without treatment. Scale bar = 100μm. **K)** Quantification of immunofluorescence staining for T_regs_ (CD3^+^CD4^+^CD25^++^) in skin of DKO*^K15^ mice as percentage of CD4^+^ T cells (left) and absolute numbers/mm^2^. Each data point represents one skin section. **L)** Representative images of SSAT activity staining in mouse skin with and without treatment. Inhibitor control = addition of diminazene aceturate to measure specific activity. Scale bar = 100μm. **M)** Quantification of mean fluorescence intensity of SSAT activity in the skin of DKO*^K15^ mice with and without treatment. **A)** and **B)** n=4 mice per group with two sections per mouse pooled, two-tailed unpaired t-test, *p<0.05. Mean ± SD. **E)** and **I)** n=8-9 mice per group combined from 3 different experiments, two-tailed unpaired t-test, *p<0.05, **p<0.01, ***p<0.001. Mean ± SD. **H)** n=8-9 mice per group form 3 different experiments, Two-way ANOVA with Sidak correction for multiple testing, *p<0.05, **p<0.01, ***p<0.001. Mean ± SD. **K)** and **M)** Data from 4 mice per group with 2-3 sections stained per mouse, two-tailed unpaired t-test, *p<0.05, **p<0.01, ***p<0.001. Mean ± SD.

DKO*^K15^ mice were treated with 20mg/kg DA (or NaCl as a vehicle control) every 24h (Fig. 5D) for 10 days. Upon treatment, CD4^+^ T cell composition in the spleen was assessed and a significant increase in T_reg_ percentage and absolute numbers as well as a decrease in the T_eff_:T_reg_ ratio in DA-treated mice compared to vehicle-treated controls (Fig.5E and F) was observed. Furthermore, spleen-derived T_regs_ and CD4^+^ T_eff_ cells were isolated and used to set up an *ex vivo* suppression assay to assess systemic T_reg_ function upon DA treatment. While the capacity of CD4^+^ T_eff_ to react to TCR stimulation and subsequent proliferation was not affected by DA treatment, T_reg_ suppressive ability was increased upon DA treatment. Similar to aSAT1 human T_regs_, T_regs_ from vehicle-treated mice did not suppress T_eff_ cell proliferation, whereas T_regs_ from DA-treated mice decreased T cell proliferation (Fig. 5G and H). We then isolated RNA from spleen-derived T cells and found a significant decrease in *SAT1* and *IL17A* expression in DA-treated T_regs_ (Fig. 5I). However, there was no significant change in CD4^+^ T_eff_ numbers or percentage as well as in the expression of *SAT1* (Extended Data Fig. 9C-E). Even though the phenotype of T_effs_ was not significantly changed upon DA treatment, a significant reduction of *IL17A* expression in these cells after DA-treatment was observed (Extended Data Fig. 9D). As DA treatment rescued the number and function of T_regs_ systemically, we also investigated whether DA treatment can reverse the phenotype in the skin. Immunofluorescence staining of the skin of DA– and NaCl-treated mice for T_regs_ (Fig. 5J) revealed that, like in the spleen, T_reg_ numbers and percentage in the skin after DA-treatment compared to the vehicle control were significantly increased (Fig. 5K), while SSAT activity was decreased (Fig. 5L and M). However, no significant histological change in the skin of DA-treated mice was seen (Extended Data Fig. 9F), most likely due to the short treatment window and the kinetics of cutaneous inflammation of DKO*^K^^15^ mice. However, an improvement in T_reg_ number and function within this short time was already apparent, which is a promising indication that inhibition of SSAT could have therapeutic benefit in T_H_17-mediated skin inflammation. Together, this shows that, the functional consequences of *SAT1* expression and SSAT function is specific to T_regs_ and its inhibition rescues T_reg_ function and number systemically and locally in the skin *in vivo*. However, its inhibition by DA either non-specifically inhibits *IL17A* expression or has an indirect effect on T_eff_ by increasing T_reg_ suppressive function, thereby potentially breaking a continuous IL17-driven inflammation loop and allowing for T_regs_ to recover.

To translate our preclinical findings into a clinical setting, frozen peripheral blood mononuclear cells (PBMCs) from psoriasis and sarcoidosis patients were used to assess T_regs_ (CD4^+^CD25^hi^CD127^lo^, Extended Data Fig. 9I) for a SAT1^high^ phenotype (CXCR6, CD69, PD-1) by flow cytometry. As expected, there was an overall reduction in the percentage of T_regs_ in patients suffering from psoriasis or sarcoidosis (Extended Data Fig. 9G). T_regs_ in the blood of these patients already had a higher expression of CXCR6 compared to healthy controls as well as a decreased expression of CD69 and no change in PD-1 (Extended Data Fig. 9H). This indicates that patients suffering from chronic inflammatory skin disease already have an increased number of CXCR6^+^ T_regs_ in circulation, which could exacerbate the phenotype once these cells migrate to the skin.

## Discussion

Here we show for the first time that human T_regs_ from inflamed skin expressing the rate-limiting enzyme in polyamine metabolism, SSAT, lose their ability to control effector T cell proliferation and switch their phenotype towards a T_H_17-like expression profile. Furthermore, we demonstrate that a T_H_17 skin microenvironment can induce this phenotype via co-receptors of keratinocytes and highlight the therapeutic potential of targeting T_regs_ to restore their function by blocking SSAT in an *in vivo* mouse model of chronic skin inflammation. Therefore, inhibiting SSAT in patients suffering from chronic skin inflammation and reducing its activity systemically is a promising, T_reg_-specific, therapeutic strategy to develop in the future.

While these data support the recent reports in mouse models that polyamine metabolism is important for initial polarization of helper T cells^10, 11^, we observed no differences or phenotypic switch in human CD4^+^ effector T cells, which are already past their initial polarization, upon specific modulation of polyamine catabolism. This indicates that established T_regs_ in humans seem to be specifically vulnerable to alterations in polyamine levels, independent of peripheral polarization from naïve T cells. This aligns with a previous report that metabolic changes can induce a T_H_1-T_reg_ phenotype in which established T_regs_ can switch their phenotype to an effector cell lineage accompanied by a downregulation of FoxP3 in an autoimmunity setting^36^. In our setting, SSAT specifically induces upregulation of T_H_17 markers, effector molecules, and loss of suppressive function, while the expression of FoxP3 remains unchanged. A similar observation has been made in the context of opioid addiction, where T_regs_ producing IFN-γ but expressing FoxP3 have been termed “fragile T_regs_” and have been found to be responsible for mediating opioid withdrawal symptoms^9^.

Our study agrees more closely with the definition of a fragile T_reg_ phenotype rather than a full switch into a different T cell lineage, indicating that the T_reg_ phenotype can potentially be rescued by the reversal of pathological characteristics of T_regs_ in various inflammatory diseases. Many chronic inflammatory conditions have a T_H_17 background, and recently it has also been shown that traditionally T_H_2-mediated diseases, such as AD, have a pronounced T_H_17 signature^21, 22^. scRNA-seq data from AD patients also showed an increase in T_H_17 signature genes among most clusters. However, using a “clean” T_H_2 environment (i.e., IL-4) *in vitro*, an induction of a SAT1^high^ phenotype in healthy T_regs_ was not observed. Along the same lines, keratinocytes have been implicated in being a disease-driver in a T_H_17-inflammatory loop^27^, with IL-6 production even being linked to polyamine metabolism in keratinocytes themselves^37^. Given the strong signal of SSAT activity in the epidermal layer of DKO*^K15^ mice, this could have an indirect effect on surrounding T cells in conjunction with direct T cell modulation via co-receptors. In addition to what has previously been reported^28, 29, 30, 31^, we also observe that keratinocytes can upregulate co-receptors to interact with T cells. This adds to existing studies that have implicated keratinocytes in directly modulating T cell responses in the skin^38, 39^. While the mouse model used in this study is driven by keratinocytes, it would be interesting to test different models of chronic skin inflammation for systemic treatment as well as having an increased topical treatment period to see if there would be any long-term histological changes. However, it cannot be excluded that *SAT1* expression in T_regs_ can also be induced by co-receptor expression of other cell types. The findings presented here further expand the knowledge on antigen-presenting-cell-independent priming of T cell responses in tissues and highlight the importance of structural cells in mediating immune responses. Linking inflamed keratinocytes in a T_H_17 microenvironment to the T_reg_ phenotype observed in this study opens additional avenues for targeting T_regs_ in chronic inflammatory skin disease.

Even though immunotherapies exist for these patients, these broadly block an effector cytokine or its signaling pathway, leading to side effects in form of opportunistic infections^6, 7^. It would be of great benefit, if therapies were developed that specifically aim to reinstate proper T_reg_ function and permanently break this inflammatory loop. Here, we identify the acetyltransferase SSAT as one such target. The inhibitor applied in this study, DA, is primarily used in veterinary medicine under the trade name Berenil to treat trypanosomiasis, but is also FDA approved for human use in protozoan infections^40^. As prolonged DA treatment is known to cause neurological side effects in animals^41^, we used a short treatment interval to demonstrate that even short-term inhibition of SSAT could potentially improve T_reg_ function *in vivo*. As improvement of T_reg_ number and function after a 10-day treatment period was already observed, developing a small molecule inhibitor for SSAT may be a novel avenue to restore T_reg_ function and attenuate T_eff_ cells^42^ in chronic inflammation. As expression of *SAT1* within the T cell compartment is specific to T_regs_, and there also was a specific downregulation in T_regs_ upon treatment, using a pan-inhibitor already indicates a T_reg_-specific effect without having to develop sophisticated T_reg_-targeting methods to boost T_reg_ function and restore immune homeostasis. Targeting polyamines could have further benefits for general health, as aging and aging-related diseases have been associated with dysregulated polyamines (reviewed in refs.^43, 44^) and a dysfunctional T_reg_ population^45, 46^, which has also been linked to TNFα, a known inducer of *SAT1* gene expression^20^ in aged skin^47^. It is interesting to speculate that SAT1^high^ T_regs_ are also involved in “inflammaging”, and that modulating polyamine metabolism would attenuate T_reg_-mediated aging symptoms and be one mechanism by which aging-associated disease could be additionally treated.

In conclusion, this data employing T_regs_ derived from human skin, demonstrates how environmental factors alter polyamine metabolism, leading to a transition in T_regs_ towards a pro-inflammatory phenotype, characterized by the production of effector cytokines and a loss of their suppressive capacity. While we characterize *SAT1*-induced dysfunctional T_regs_, the exact molecular mechanism by which this phenotype is induced remains unclear and was mainly limited by cell availability from human tissue. Still, this study demonstrates how targeting this specific enzyme could be of therapeutic benefit in chronic inflammatory skin diseases by restoring T_reg_ function and breaking the continued inflammatory loop.

## Methods

### Patient cohort and recruitment

Untreated patients suffering from psoriasis and sarcoidosis were recruited via the Department of Dermatology at the Medical University of Vienna (Supplementary Table 1) and donated 56mL of blood. All patients participated in this study voluntarily and have given fully informed written consent. This has been approved by the ethics committee of the Medical University of Vienna (ECS 1503/2020).

### Healthy human skin and blood samples

Healthy donor blood was obtained from healthy volunteers. Skin was obtained from elective surgeries from the Department of Plastic and Reconstructive Surgery at the Medical University of Vienna. The research is in line with national law and approved by the Ethics Committee of the Medical University of Vienna (ECS 1890/2018).

### Mouse model

All mouse experiments were performed in accordance with local and institutional regulations/licenses and comply with national and European guidelines (EU Directive 2010/63/EU for animal experiments). Mouse work was approved under Wagner E. 171/18. Male and female JunB^f/f^;c-Jun^f/f^;K15-CrePR adult mice^33^ were injected with Mifepristone (2mg/100μL) at 8 weeks of age, for 5 consecutive days. At 10 weeks of age, mice were treated with 20mg/kg diminazene aceturate (DA). DA was dissolved in NaCl and 100-200μL were injected intraperitoneally every 24h for 9 days. Control mice were injected with 100-200μL NaCl every 24h for 9 days. Mice were sacrificed on day 10 post-treatment and different tissues including spleen, ear, and backskin were harvested. Skin samples were snap frozen in OCT Compound Cryostat Embedding medium (Scigen, Cat: 4586) and stored at –80°C. Spleens were manually digested and cells were utilized directly.

### Immunofluorescence staining of skin sections

Skin biopsies were snap frozen in OCT and stored at –80°C. Sections were cut in a cryostat chamber to a thickness of 5-10μm and fixed in acetone and stored at –20°C. Slides were temperature compensated for 10 minutes PBS at room temperature prior to use. Tissue sections were then blocked for 20 minutes in 2% Bovine Serum Albumin (BSA) (Sigma, A2153) in PBS containing 2% of the appropriate serum, depending on antibody isotype. For FoxP3 staining, blocking solution also contains 0.3% TWEEN® 20 (Sigma, Cat: P1379). Intracellular and intranuclear markers were stained first for 2h at room temperature. For SSAT stainings, a second step mouse anti-rabbit AF488 (SantaCruz, Cat: sc-2359, 1:200) was performed for 1h at room temperature. Cell surface markers are stained overnight at 4°C. Tissue was then stained for 5 minutes with DAPI (Sigma, Cat: D9542-1MG) and mounted in PermaFluor™ Epredia™ Lab Vision™ aqueous mounting medium (ePredia, Cat: TA-030-FM), covered with a coverslip and stored at 4°C until acquisition. Tissue staining was acquired with the TissueFAXS system (TissueGnostics, Austria) and analyzed using the corresponding TissueQuest software (TissueGnostics, Austria). Antibodies used for human staining were anti-FoxP3 PE (Biolegend, Cat: 320108, 1:100), anti-CD3 APC (BD, Cat: 345767, 1:100), anti-CD4 FITC (BD, Cat: 345768, 1:50), anti-SAT1 (abcam, Cat: ab105220, 1:50), and Mouse anti-rabbit IgG AF488 (SantaCruz, Cat: sc-2359, 1:200). For mouse samples, sections were stained for 3h at room temperature with anti-CD3 PE-Dazzle594 (Biolegend, Cat: 100245, 1:50), anti-CD4 PE (Biolegend, Cat: 100408, 1:20), and anti-CD25 APC (Biolegend, Cat: 102012, 1:100). Respective isotypes were diluted to the same concentration and stained in parallel to antigen-specific antibodies.

### Immune cell isolation from PBMCs

Whole blood was collected in heparin tubes and PBMCs were isolated by Ficoll-Paque (GE Healthcare, Cat: 17-1440-03) gradient centrifugation. Isolated PBMCs were either used immediately for further isolation of immune cell subsets, or frozen in Fetal Bovine Serum (FBS, Gibco, Cat: 10500064) with 10% DMSO (Sigma, Cat: 317275-500ML) at –80°C. T_regs_ were isolated from fresh PBMCs by magnetic separation using the CD4^+^CD25^+^CD127^dim/-^ human T_reg_ isolation kit (Miltenyi Biotec, Cat: 130-094-775). T_regs_ were then used directly for analysis or assays.

### T cell expansion from human skin biopsies

Skin explants were prepared as described by Clark et al^47^. Briefly, skin biopsies were cut into small places and placed onto a collagen-coated (Collagen G, Sigma Aldrich, Cat: L7213, 1:40 in PBS) foam matrix. Matrices were placed into ImmunoCult™-XF T Cell Expansion Medium (StemCell, Cat: 10981) supplemented with rhIL-2 (200IU; Peprotech, Cat: 200-02), rhIL-7 (10ng/mL; Peprotech, Cat: 200-07), and rhIL-15 (10ng/mL; Peprotech, Cat: 200-15) and 1% Penicillin-Streptomycin (PS, Gibco, Cat: 15140122). Media with fresh cytokines was exchanged every 3-4 days. Expanded T cells were harvested after 14-16 days of culture. T_regs_ were isolated magnetically (see above).

### Sample preparation for metabolomics

T_regs_ were isolated from fresh PBMCs as described above. All solutions were pre-cooled, and the temperature of the sample is kept at 4°C to ensure retention of metabolites. Isolated cells were counted and an equal amount of T_eff_ (CD4^+^CD25^low^CD127^hi^) and T_regs_ were pelleted and snap frozen in liquid nitrogen. Samples were stored at –80°C until processing for LC-MS. Metabolite extraction and targeted analysis for polyamines and amino acids was performed by the Proteomics-Metabolomics facility at CeMM Research Center for Molecular Medicine of the Austrian Academy of Sciences.

### Nucleofection of human primary T cells

Cells isolated from skin or blood were seeded at 1×10^6^ cells/mL in ImmunoCult™-XF T Cell Expansion Medium and activated with ImmunoCult™ CD3/CD28 Human T cell activator (5μL/mL T_regs_, 25μL/mL T_eff_; StemCell, Cat: 10971) for 24h. Before nucleofection, cells were washed once with PBS and resuspended in T cell nucleofection solution (Lonza, Cat: VPA-1002) containing plasmids of choice and nucleofected using the T-023 program on the Amaxa IIb nucleofector. Cells were then immediately transferred into pre-warmed in Roswell Park Memorial Institute (RPMI)-1640 media (Gibco, Cat: 52400-025) + 10% FBS + 1% PS and incubated 48h before analysis.

### CRISPRa of SAT1 in primary human T cells

sgRNA sequences for SAT1 were taken from the Calabrese A genome-wide CRISPRa library (Addgene, Cat: 92379) and cloned into the pLKO5.sgRNA.EFS.GFP (Addgene, #57822) backbone. Cells isolated from skin were seeded at 1×10^6^ cells/mL RPMI-1640 media (Gibco, Cat: 52400-025) + 10% FBS + 1% PS + rhIL-2 and activated with ImmunoCult™ CD3/CD28 Human T cell activator for 24h. Cells were nucleofected with the vectors containing sgRNA (5’-GGGCGCCCGTGTGAACCCGG-3’ and 5’-GTGGGACCTCTCATCCAATG-3’; 1.5μg each) and a dCas9-VPR plasmid (pLenti-dCas9-VPR-blast, Addgene, #96917; 1μg) as described above, keeping the amount of DNA between non-targeting control and targeting guides constant. Cells were then immediately transferred into pre-warmed RPMI-1640 + 10% FBS + 1% PS and incubated at least 48h before analysis.

### Flow cytometry and fluorescence-activated cell sorting (FACS)

Surface markers were stained in PBS for 30min on 4°C and then washed with FACS buffer (2mM EDTA (ThermoFisher, Cat: 15575020), 1% BSA in PBS). For intracellular markers and staining without sorting, cells were fixed for 20 minutes at room temperature using a Fixation Buffer (Biolegend, Cat: 420801) followed by permeabilization (Biolegend, Cat: 421002) and 30 minutes intracellular antibody staining at room temperature. For staining of transcription factors, the eBioscience FoxP3/Transcription Factor staining buffer set (Invitrogen, Cat: 00-5523-00) was used according to manufacturer’s instructions. Data were acquired on the spectral analyzer Aurora (Cytek®) with a 5-laser setup. For cell sorting, BD FACS Aria II was used. Acquired data was analyzed using FlowJo v.10.8.

For experiments assessing the SAT1^high^ phenotype, cells were surface stained with ZombieUV (Biolegend, Cat: 423107, 1:1000), anti-CD4 BUV536 (BD, Cat: 750979, 1:400), anti-CD127 PacificBlue (Biolegend, Cat: 351305, 1:200), anti-ICOS BV480 (BD, Cat: 746248, 1:300), anti-PD-1 BV510 (Biolegend, Cat: 329932, 1:200), anti-CD28 BV650 (Biolegend, Cat: 302946, 1:100), anti-CD25 BV711 (BD, Cat: 563159, 1:50), anti-CD3 PE-Dazzle594 (Biolegend, Cat: 300450, 1:200), anti-CD69 PE-Cy5 (Biolegend, Cat: 310907, 1:100), anti-CXCR6 PE-Cy7 (Biolegend, Cat: 356011, 1:200), anti-CD27 AF700 (Biolegend, Cat: 356416, 1:200), and anti-TIGIT APC-Fire750 (Biolegend, Cat: 372708, 1:100). Cytokines were stained with anti-IFN-γ (Biolegend, Cat: 502542, 1:40), anti-IL-10 PerCP-Cy5.5 (Biolegend, Cat: 501418, 1:50), anti-IL-2 PE (Biolegend, Cat: 340450, 1:50), anti-IL-17A APC (eBioscience, Cat: 17-7179-41, 1:50). For experiments assessing transcription factor expression, cells were pre-stained with ZombieNIR (Biolegend, Cat: 423105, 1:100), sorted for GFP^+^ cells and then stained with anti-FoxP3 BV421 (Biolegend, Cat: 320124, 1:20), anti-T-bet BV605 (Biolegend, Cat: 644817, 1:50), anti-RUNX3 PE (BD, Cat: 564814, 1:50), anti-Helios PE-Dazzle594 (Biolegend, Cat: 137231, 1:20), anti-ROR-γt APC (Invitrogen, Cat: 12-6988-82, 1:50), anti-pSTAT3 (CellSignaling, Cat: 4324S, 1:100). Keratinocytes were stained with ZombieUV (Biolegend, Cat: 423107, 1:2000), anti-HVEM BUV737 (BD, Cat: 748637, 1:200), anti-CD86 BV605 (Biolegend, Cat: 374213, 1:100), anti-CD80 BV711 (Biolegend, Cat: 305236, 1:100), anti-PD-L1 BV785 (Biolegend, Cat: 329736, 1:100), anti-CD112 PerCP-Cy5.5 (Biolegend, Cat: 337415, 1:200), anti-HLA-DR PE-Cy5 (Biolegend, Cat: 555813, 1:100), anti-4-1BBL PE-Cy7 (Biolegend, Cat: 311511, 1:300), and anti-CD155 AF647 (Biolegend, Cat: 337621, 1:400).

### Sorting patient PBMCs for scRNAseq

PBMCs from psoriasis patients (n=4) were thawed into 5mL RPMI and washed once with ice cold PBS. Cells were then stained with eF780 viability dye (eBioscience, Cat: 65-0865-14, 1:1000), anti-CD4 PE-Cy7 (Biolegend, Cat: 317414, 1:200), anti-CD45 PE-TexasRed (BD, Cat: 562279, 1:200), anti-CD69 PE (BD, Cat: 555311, 1:20), anti-CD3 BV421 (Biolegend, Cat: 3004341, 1:100), and anti-CD8 APC (BD, Cat: 5553690, 1:50). Each patient was also stained with one of TotalSeq™-C0251/2/3/4 Antibody (Biolegend, Cat: 394661/3/5/7, 1:100) for sample multiplexing. Cells were stained for 30min at 4°C in the dark. After staining, cells were washed with FACS buffer and resuspended for sorting. 7500 live CD45^+^ cells per patient were sorted and pooled for library preparation and sequencing. Libraries were prepared using the Chromium Controller and the Chromium Next GEM Single Cell 5’ Kit v2 (10× Genomics, Cat: PN-1000263) according to manufacturer’s instruction. Libraries were sequenced by the Biomedical Sequencing Facility at the CeMM Research Center for Molecular Medicine of the Austrian Academy of Sciences, using the NoveSeq6000 platform. For data processing and analysis see Bioinformatic analysis of scRNA-seq data.

### T cell suppression assay

T_regs_ and CD4^+^ T_eff_ cells were isolated as described above. T_eff_ were stained with CellTrace Violet™ (ThermoFisher, Cat: C34557) according to manufacturer’s instructions to a working concentration of 1μM. An aliquot of T_regs_ was stained for FoxP3 expression to ensure a pure population of cells. T_regs_ were serial diluted into a 96-well plate containing RPMI-1640 + 10% FBS + 1% PS and an equal amount of T_eff_ was added to each well containing T_regs_ to create different ratios of T_eff_:T_reg_. Control wells were seeded using T_eff_ with and without stimulation. All wells, except no-stimulation control wells, were activated with ImmunoCult™ CD3/CD28 Human T cell activator (25μL/mL) and left to incubate for 5 days. Cells were harvested and stained for 30min at 4°C with the following antibodies: anti-CD25 BV785 (Biolegend, Cat: 356139, 1:50), anti-CD127 PE (Biolegend, Cat: 351304, 1:100) anti-CD4 PE-Cy7 (Biolegend, Cat: 317414, 1:200), and eFluor viability dye (eBioscience, Cat: 65-0865-14, 1:1000). For mouse cells, spleens were manually dissociated and filtered through a 70μM cell strainer (Falcon, Cat: 352350). T_regs_ and CD4^+^ T_eff_ were then isolated using the CD4^+^CD25^+^ Regulatory T Cell Isolation Kit, mouse (Miltenyi Biotec, Cat: 130-091-041) according to manufacturer’s instructions. Cells were activated using Dynabeads™ Mouse T-Activator CD3/CD28 for T-Cell Expansion and Activation (ThermoFisher, Cat: 11456D) according to manufacturer’s instructions. Co-culture of T_regs_ and CD4^+^ T_eff_ was set up identically to the human setup described above. After 5 days of incubation, cells were harvested and stained for 30min at 4°C with anti-CD25 APC (Biolegend, Cat: 102012, 1:50), and ZombieUV (Biolegend, Cat: 423108, 1:1000). Cell proliferation was assessed by flow cytometry and acquired data was analyzed using FlowJo v.10.8.

### RT-PCR

RNA was isolated using the RNeasy Mini spin kit (Qiagen, Cat: 74106) or RNeasy Micro kit (Qiagen, Cat: 74004), depending on cell number, according to manufacturer’s instruction. RNA quality was assessed using agarose gel electrophoresis (1.5% w/v). Samples were retrotranscribed to cDNA using the Biozym cDNA synthesis kit (Biozym, Cat: 331470X/L) according to manufacturer’s protocol. Primers were designed using the NCBI primer designing tool. Primers were designed following requirements for RT-PCR. Self– and cross reactivity of selected primers was simulated with Beacon Designer. Selected primers were also run through a BLAST search to confirm specificity to a single gene. Primer pairs (forward, reverse) used for human samples were *RSP18* 5’-GTTCCAGCATATTTTGCGAGT-3’, 5’-GTCAATGTCTGCTTTCCTCAAC-3’; *SAT1* 5’-AGAAGATGGTTTTGGAGAGC-3’, 5’-CATCACGAAGAAGTCCTCAA-3’; *PAOX* 5’-GGAGAGTGACAGGAAACCC-3’, 5’-CGTGGAGTAAAACGTGCG-3’; *SRM* 5’-CGCAGTAAGACCTATGGC-3’, 5’-ATGATCAGCACCTTTCGC-3’; *ARG2* 5’-CTGGCTTGATGAAAAGGCTCTCC-3’, 5’-TGAGCGTGGATTCACTATCAGGT-3’; *ODC1* 5’-CCAAAGCAGTCTGTCGTCTCAG-3’, 5’-CAGAGATTGCCTGCACGAAGGT-3’; *AMD1* 5’-ACCACCCTCTTGCTGAAAGCAC-3’, 5’-CCCTTGGTGAGAAGGCTTCATG-3’; *SMOX* 5’-ACGGAGATGCTGCGTCAGTTCA-3’, 5’-CCTGCGTGTATGAATAGGAGCC-3’; *IL17A* 5’-CCATAGTGAAGGCAGGAATC-3’, 5’-TCCTCATTGCGGTGGA-3’; *IL10* 5’-TCCTGCCCTTAGGGTTAC-3’, 5’-GCCTTTCTCTTGGAGCTTAT-3’; for mouse samples primer pairs used were the following: *RPL13a* 5’-AGCCTACCAGAAAGTTTGCTTAC-3’, 5’-GCTTCTTCTTCCGATAGTGCATC-3’; *SAT1* 5’-ACCGTCTTGCCACTTCTTAG-3’, 5’-CCAACAATGCTATGTCCTTCA-3’; *IL17A* 5’-GGAGAGCTTCATCTGTGTCTC-3’, 5’-CTTCACATTCTGGAGGAAGTC-3’; *IL10* 5’-TTTACCTGGTAGAAGTGATGC-3’, 5’-ACCTTGGTCTTGGAGCTTAT-3’. All PCR reactions were performed using the PowerUp SYBR Green Master Mix (ThermoFisher, Cat: A25741) on the StepOnePlus Real-Time PCR system with StepOne Software v2.3 (Applied Biosystems, Thermo Fisher Scientific) according to manufacturer’s instructions. CT values were used to calculate the relative expression level (comparative CT or ddCT-method) with RSP18 (human) or RPL13a (mouse) as a housekeeping gene. Relative expression levels were expressed as fold change relative to the control.

### T_reg_-keratinocyte co-culture

NHEK-svTERT3-5 (Evercyte, Cat: CLHT-011-0026-5) keratinocytes were cultured in KGM-2 media (Promocell, Cat: C-20111). Keratinocytes were plated at 2.5×10^5^/mL in a 48-well plate and left to attach overnight. The next day, keratinocytes were stimulated with IFN-γ (50g/mL; Peprotech, Cat: 300-2), IL-1β (10ng/mL; Peprotech, 200-01B), and IL-17A (100ng/mL; Peprotech, Cat: 200-17) overnight. T cells were isolated from whole blood as described above and incubated overnight with ImmunoCult™ CD3/CD28 Human T cell activator (5μL/mL T_regs_, 25μL/mL T_eff_) in ImmunoCult™-XF T Cell Expansion Medium. To set up the co-culture, T_H_17-stimulating media was removed from keratinocytes and fresh KGM-2 was added. T cells were added at 1.25×10^5^/mL in ImmunoCult™-XF T Cell Expansion Medium for a final 1:1 mix of KGM-2:ImmunoCult™. The co-culture was incubated for 5 days before harvest. T cells were washed off the keratinocyte layer using cold PBS and either used for flow cytometry or RNA extraction. Keratinocytes were lifted off the plate using Trypsin/EDTA (Lonza, Cat: LONCC-5012) for 5min at 37°C. Trypsinization was stopped using trypsin neutralizing solution (Lonza, Cat: LONCC-5002) and PBS. Cells were then washed and used for flow cytometry.

### Histochemical enzyme activity assay for SSAT activity in situ

Skin biopsies were snap frozen in OCT (Scigen, Cat: 4586) and stored at –80°C. Sections were cut in a cryostat chamber to a thickness of 5μm and transferred directly to –80°C without fixative to prevent interference with enzyme activity. Sections were briefly thawed at room temperature and then stained with the enzyme reaction mix for 5min. Slides were stained and imaged individually and consecutively. The reaction mix contained the following: Amplite® Fluorimetric Coenzyme A Quantitation Kit Green Fluorescence (1X; AAT Bioquest, Cat: 15270) was used to detect acetylation, spermine (2mM; Merck, Cat: S4264) and spermidine (2mM; Merck, Cat: S0266) as enzyme specific substrates, acetyl coA (0.5mM; Merck, Cat: A2181) as a co-factor together with DAPI (1X; Sigma, Cat: D9542-1MG). All reagents were added to a buffered solution containing water and Tris-HCl (100mM, pH 7.5) and polyvinyl alcohol (PVA, 18%, pH 7.5) to have a final concentration of 50mM and 100μL were used per slide. As negative control, the SSAT inhibitor Diminazene aceturate (20μM; Merck, Cat: D7770) was added to the assay mixture. Enzyme reactions were carried out protected from light. Immediately after 5min incubation, a coverslip was mounted and slides were imaged within 10min. Images were acquired using the TissueFAXS system, using the 514nm channel to acquire the enzyme activity staining and DAPI for nuclear staining. Analysis was performed with the TissueQuest software. Enzyme activity was calculated using mean fluorescence intensity (MFI) of the enzyme activity signal relative to the mean fluorescence intensity of the enzyme stain containing the inhibitor. Relative signal was calculated by subtracting the background signal in the negative control, inhibitor-treated, slides.

## Bioinformatic analysis of scRNAseq data

### Single-cell RNA-seq data processing

The raw sequencing data for each sample was processed with cellranger v6.0.1 count (Psoriasis/AD skin and PBMC, Ulcerative colitis data) or multi (Sarcoidosis skin and PBMC data) using the 10x GRCh38 v3.0.0 reference. The processed samples were then combined using cellranger aggr. Doublet cells were detected with doubletdetection v4.2 (http://doi.org/10.5281/zenodo.2678041) and removed. In total this resulted in the generation of data for 103,334 cells (Ulcerative colitis), 384,215 cells (AD skin), 45,977 cells (Psoriasis/Sarcoidosis PBMC) and 422,087 cells (Psoriasis/Sarcoidosis skin) with 33,538 annotated genes.

### Single-cell RNA-seq data quality control

Each of the analyzed datasets was quality controlled by applying thresholds on the number of detected genes, the percentage of mitochondrial RNA as well as the percentage of ribosomal RNA per cell. In brief, cells were removed if their percentage of mitochondrial RNA was higher than 15%, their number of detected genes was lower than 750 (Psoriasis skin and PBMC, Sarcoidosis), 400 (Ulcerative colitis colon), 500 (AD skin) or higher than 5000 and their percentage of ribosomal RNA was lower than 5%. We aimed for a number of retained genes per dataset of approximately 20000. Therefore, genes expressed in less than 50 (AD skin, Psoriasis/Sarcoidosis PBMC), 100 (Psoriasis/Sarcoidosis skin) and 10 cells (Ulcerative colitis colon) were removed. This quality control resulted in final dataset sizes of 79,378 cells and 18,436 genes (Ulcerative colitis), 318,976 cells and 21,750 genes (AD skin), 40,148 cells and 15,998 genes (Psoriasis/Sarcoidosis PBMC) and 332,546 cells and 20,912 genes (Psoriasis/Sarcoidosis skin).

### Identification of regulatory T cells

The identification of T_regs_ was done in two iterations of data integration and clustering. In brief, each dataset was first integrated the quality-controlled data with scVI v0.15.4^48^ (n_layers = 2, n_latent = 30, gene_likelihood = ‘nb’, train_size = 1) using only the top 4000 highly variable genes and the Sample ID as batch key. We then computed a k-Nearest Neighbor graph on the resulting scVI embedding and used this to subsequently cluster the graph with the Leiden algorithm (implemented in the leidenalg package v0.10.2 (https://github.com/vtraag/leidenalg) and executed via scanpy v1.9.7 ^49^ with resolutions 0.1 (Psoriasis/Sarcoidosis skin), 0.4 (Ulcerative colitis colon), 0.25 (AD skin) and 0.3 (Psoriasis/Sarcoidosis PBMC). We then regarded those clusters with the highest CD3D expression as T cells, subsetted the data accordingly and again integrated with scVI, this time, using all available genes, computed the k-Nearest Neighbor graph on the scVI embedding and subsequently clustered using the Leiden algorithm with resolutions 0.4 (Psoriasis/Sarcoidosis skin), 1.0 (Ulcerative colitis colon), 0.3 (AD skin) and 0.5 (Psoriasis/Sarcoidosis PBMC). Clusters with the highest FOXP3 expression were then regarded as regulatory T-cells. All data was visualized using Uniform Manifold Approximation and Projection (UMAP) v0.5.3 ^50^.

### Differential cell abundance analysis

Differential cell abundance in disease versus healthy condition was assessed with the milopy package v0.1.0^19^ (underlying R version 4.0.5 interfaced with rpy2 v3.5.15). In brief, cell neighborhoods were computed based on the UMAP space computed from the previously generated scVI embedding. Cell abundance differences were then assessed based on cell counts per Sample ID in each neighborhood.

### Determining SAT1 expression state of regulatory T cells

Psoriasis/Sarcoidosis skin T_regs_ were subdivided into one of two states based on their SAT1 expression and neighboring cells. In brief, we first integrated the T_reg_ subset of the Psoriasis/Sarcoidosis T-cells with scVI. We then computed the log-normalized expression (target sum = 10,000 reads per cell) and used the median SAT1 expression as a threshold to divide the cells into SAT1^hi^ and SAT1^lo^ T_regs_. To make sure we also incorporate cells that may be SAT1^hi^ but where SAT1 was not detected we subsequently clustered the data using the Leiden algorithm on the k-Nearest Neighbor graph computed from the scVI embedding with a resolution of 5 and assigned all cells in each cluster to a given state based on a majority vote.

### Cell-cell communication analysis

Cell-cell communication between cell types was assessed with CellChat v2.1^51^ (underlying R version 4.3.0). In brief, we first clustered the integrated Psoriasis/Sarcoidosis skin data using the Leiden algorithm with a resolution of 0.75. Subsequently, we used celltypist v1.6.2 ^52^ with the ‘Adult_Human_Skin.pkl’ model to annotate the resulting clusters with their corresponding cell type (T_regs_ were further split into SAT1^hi^ and SAT1^lo^ state based on their SAT1 expression and cell neighborhood; see Determining SAT1 expression state of regulatory T cells). We then used this annotation, together with the log-normalized expression data (normalized to a target sum of 10,000 reads per cell) as input for CellChat.

### Gene set scores

The gene set scores were computed analogous to Triosh et al.^53^ on the basis of the log-normalized expression (target sum = 10,000 reads per cell) of a given list of genes corresponding to the respective score to be computed (T_reg_ fragility score: IL7R, CD69, LGALS1, CSF2, TBX21, MTOR, IL6, GAPDH, IFNG, CCR5, CXCR3, ITGAE, RORC, IL13, CD28, CCR2, CCL4, KLRG1, HIF1A, BATF, PDCD1, IL17A, RORA, STAT3, IL22, ICOS, CXCR6, CD58, CCL5, CD27, IL2; T_H_17 score: IL22, IL17A, IL32, IL17F, RORC, IL21, STAT3, LTA, IFNG, CCR6, CXCR3, CCL5, RORA, BATF; T_H_2 score: IL6ST, GATA3, IL6, STAT6, IGFBP7, STAT5B, IL4R, PPARG, VDR, IL13, IL1RL1, IL4, STAT5A). In brief, we grouped all genes into 25 bins of equal size based on their average expression over all cells. For each of the score genes we then randomly chose 100 genes without replacement from the same bin the gene is contained in and added these 100 genes to a set of control genes. We then computed the average expression of score genes and control genes per cell. The final fragility score is then given by subtracting the average control gene expression from the average score gene expression.

### Gene ranking of enriched Milo neighborhoods in **T**_regs_

Gene ranks of T_regs_ in enriched Milo neighborhoods versus all other T_regs_ were computed with the Wilcoxon rank sum test implemented in the rank_genes_groups function of scanpy v1.9.7. In brief, we computed differentially abundant neighborhoods of T_regs_ as described above. We then extracted all T_regs_ assigned to neighborhoods significantly enriched in disease (padj < 0.1, log2-Foldchange > 0) and assigned them to one group. Subsequently, we computed gene ranks via a Wilcoxon rank sum test based on the described group of enriched T_regs_ versus all other T_regs_ in the dataset. The top 25 genes (i.e., those with the highest ranks/score) were then plotted.

### Differential gene expression analysis

Differential gene expression was assessed with MAST v1.26.0^54^ executed via the FindMarkerGenes function implemented in Seurat v4.3.0.1 ^55^ (underlying R version 4.3.0). In brief, we exported the sparse raw count data to Matrix Market format and the associated metadata as tab-separated files and subsequently imported it into Seurat. We then log-normalized the data (normalization.method = ‘LogNormalize’, scale.factor = 10000) and used the FindMarkerGenes function to compute expression differences between disease and healthy conditions or SAT1 expression state (test.use = ‘MAST’, min.pct = 0.1, logfc.threshold = 0). Finally, we regarded genes with an adjusted p-value < 0.001 and an absolute log2-fold change > 0.25 as differentially expressed.

### Gene set enrichment analysis

Differentially expressed genes were subject to a gene set enrichment analysis against the MSigDB Hallmark 2020 gene set collection as provided by the Enrichr API^56^. In brief, we extracted differentially expressed genes from the MAST results (padj < 0.001, abs(log2FC) > 0.25) and split them into up– and downregulated genes. We then used the gseapy package v1.1.1 ^57^ to submit these lists to the Enrichr API retaining only those gene sets with an adjusted p-Value < 0.05.

### Data and code availability

The raw sequence data generated in this study will be deposited on GEO and will be <u>made available upon publication. Downloaded data can be</u> found at the following accessions E-MTAB-8142 (Psoriasis and AD skin), GSE125527 (Ulcerative colitis colon), EGAS00001006970 (Sarcoidosis skin and PBMC). Additionally, we also downloaded healthy donor data from GSE184320 (skin and PBMC). All analysis code for the results in this paper will be made available upon publication of the manuscript.

### Statistical analysis

Statistical analysis was performed in GraphPad Prism 9. Statistical significance was assessed by a two-tailed paired t-test (for paired samples), two-tailed unpaired t-test (for unpaired samples), or Two-Way ANOVA with correction for multiple comparisons. Only p values of 0.05 or lower were considered statistically significant (ns [not significant], p > 0.05; *, p < 0.05; **, p < 0.01; ***, p < 0.001)

## Acknowledgements

We thank the Biomedical Sequencing Facility at the CeMM Research Center for Molecular Medicine of the Austrian Academy of Sciences for sequencing the psoriasis blood samples. This work was supported by a grant from the LEO Foundation (GS, LF-OC-21-000806), an Austrian Science Fund grant (GS, FWF PAT 8019123) and a Vienna Science and Technology Fund project (GS, NXT22-005). AH is supported by FWF Sonderforschungsbereich F83. This work was supported by the Vienna Science and Technology Fund (WWTF) through project VRG15-005 granted to J.M. The Wagner laboratory is supported by the ERC (AdG 2016-741888-CSI-Fun), a H2020 – MSCA grant (ITN 2019-859860-CANCERPREV) and the Medical University of Vienna.

## Author contributions

T.N. designed and performed experiments, analyzed data, made figures and wrote the manuscript, D.M. analyzed scRNA-seq data, made figures, and wrote the manuscript, K.K. and P.T. performed mouse experiments, L.K. procured healthy blood samples, A.R. procured psoriasis and sarcoidosis blood samples, C.F. provided healthy skin samples, N.M. processed and analyzed patient metabolomics data, E.P. and A.H. provided support in setting up enzyme histochemistry staining, D.S. and A.P.K provided plasmids and input for CRISPRa experiments, E.W. provided mice and comments and discussion, J.M. provided comments and discussion, G.S. supervised the project and edited the manuscript.

