## Extended Data and Supplemental Tables for "Spermidine/spermine N1-acetyltransferase controls tissue-specific regulatory T cell function in chronic inflammation"

**Extended Data Figure 1**: Bioinformatic processing of publicly available scRNA-seq data.

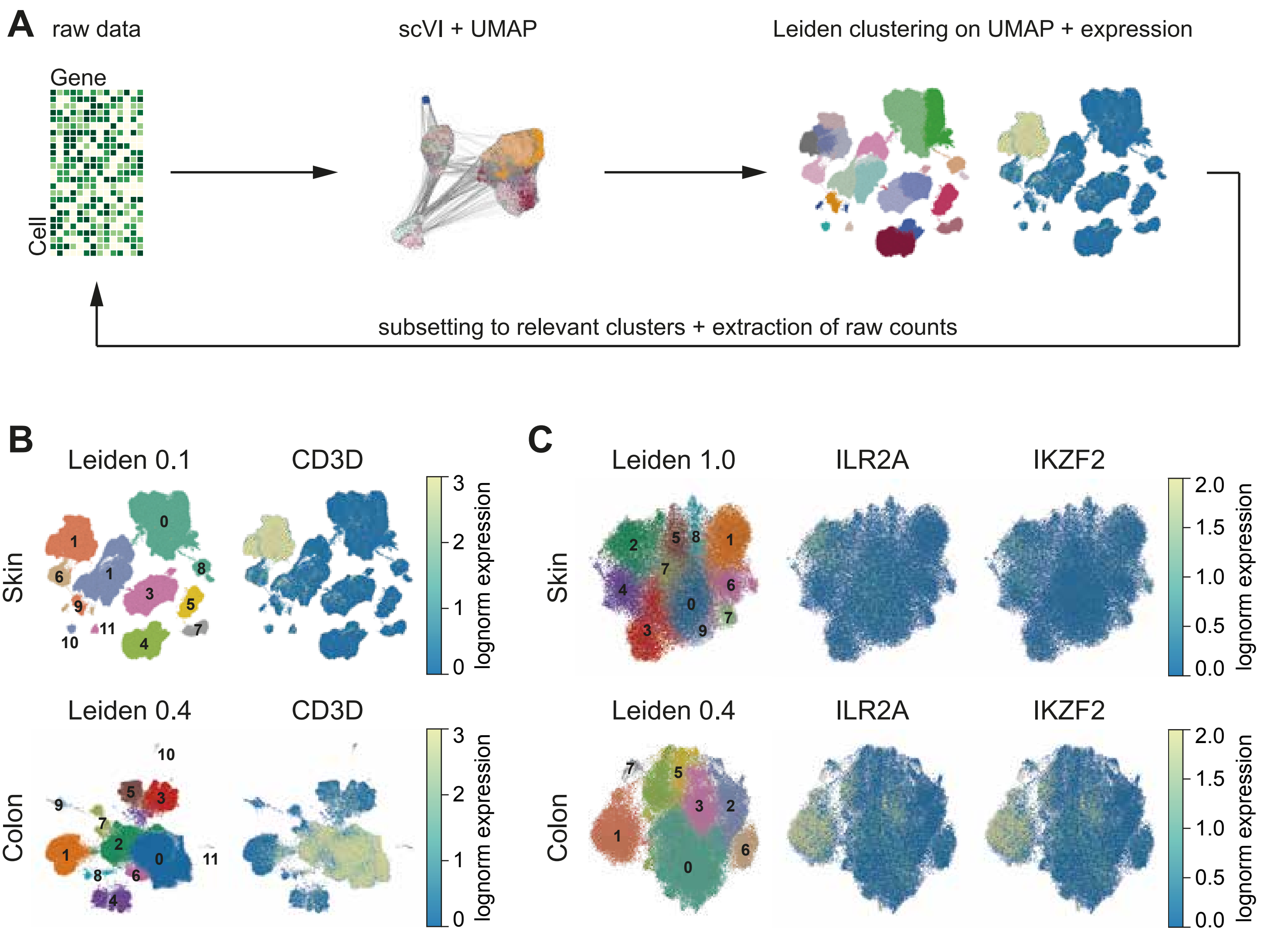

**Extended Data Figure 1: Bioinformatic processing of publicly available scRNAseq data.** **A)** Visualization of data processing, integration, and unsupervised clustering. **B)** Leiden clustering of skin cells with CD3D skin (top) and colon (bottom). **C)** Leiden clustering of skin (top) and colon (bottom) T cells showing expression of IL2RA and IKZF2.

**Extended Data Figure 2**: *SAT1* expression is specific to $T_{regs}$ in the skin.

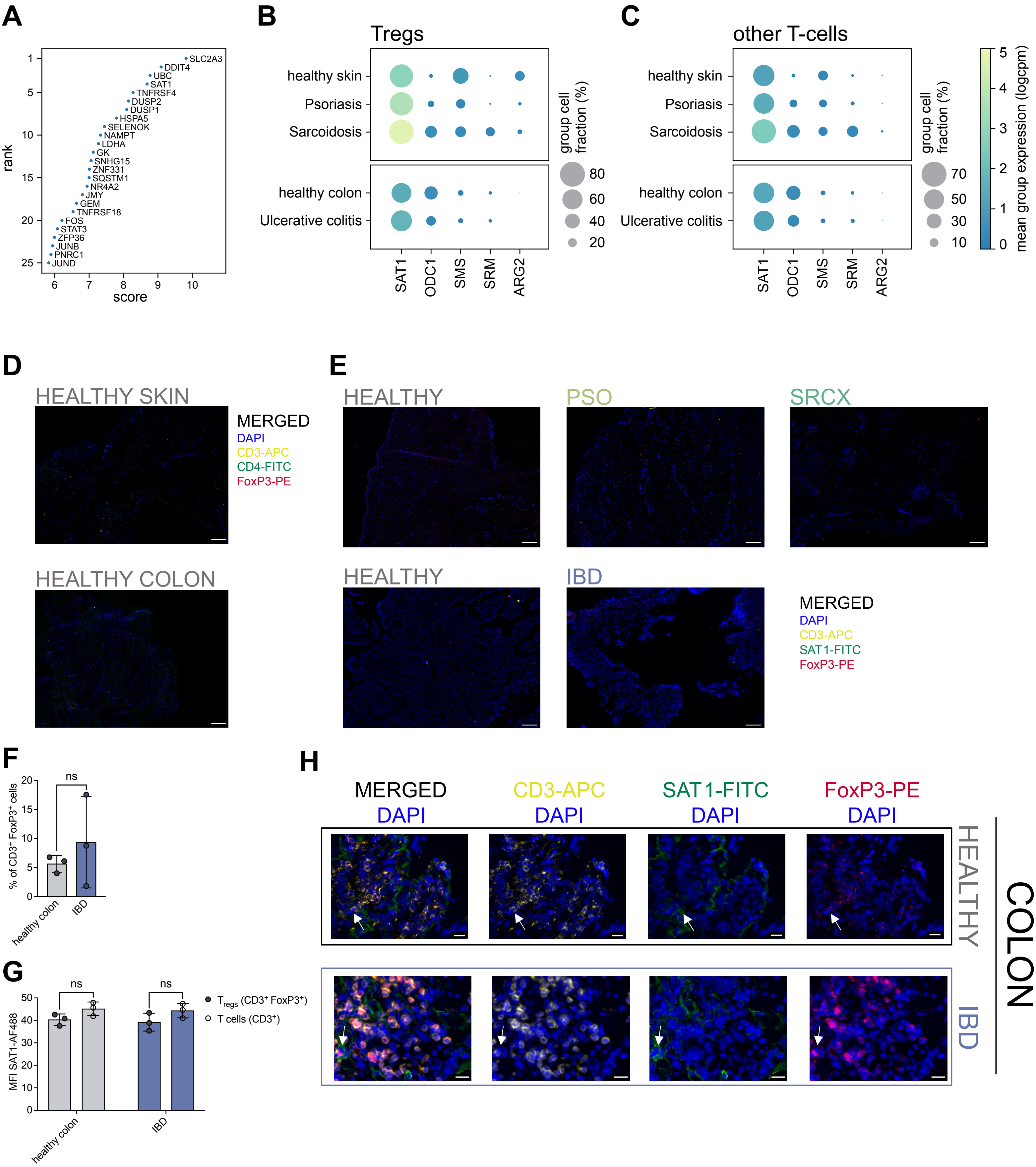

**Extended Data Figure 2: *SAT1* expression is specific to $T_{regs}$ in the skin.**

**A)** Top 25 ranked genes of overrepresented $T_{reg}$ neighborhoods according to overrepresentation analysis with Milo. Ranking based on Wilcoxon rank sum test **B)** and **C)** Dotplot of expression of genes involved in polyamine metabolism of **(B)** $T_{regs}$ and **(C)** non-$T_{regs}$ in the skin and colon. **D)** and **E)** Isotype staining of skin and colon samples for **(D)** T cells and $T_{regs}$ and **(E)** SSAT expression in T cells and $T_{regs}$. Scale bar = 200µm. **F)** Quantification of immunofluorescence staining of $SSAT^+$ $T_{regs}$ ($CD3^+$ $FoxP3^+$) in the colon. **G)** Mean fluorescence intensity of SSAT-AF488 immunofluorescence staining in $T_{regs}$ ($CD3^+$ $FoxP3^+$) and non-$T_{reg}$ T cells ($CD3^+$) in the colon. **H)** Representative images of SSAT staining in T cells in the colon. Scale bar = 20µm. Each dot represents one patient or healthy donor. Two-way ANOVA with Holm-Sidak multiple-testing correction, ns $p>0.05$. Mean ± SD.

**Extended Data Figure 3: Blood $T_{regs}$ from patient suffering from chronic skin disease express *SAT1* and have a reduced polyamine content.**

**A)** Leiden clustering of PBMCs from healthy donors (n=3) and patients (n=4 psoriasis; n=14 sarcoidosis) with inflammatory skin disease. **B)** UMAP of PBMCs showing expression of CD3D in healthy and patient PBMCs. **C)** Leiden clustering of T cells from healthy and patients with inflammatory skin disease. **D)** and **E)** UMAP of healthy and patient PBMC-derived T cells showing expression of (D) FoxP3 and (E) SAT1. **F)** Flow cytometry analysis of $T_{regs}$ (top) and $T_{eff}$ (bottom) after magnetic isolation based on CD127 and CD25. **G)** Relative abundance of intracellular spermidine and spermine in $T_{regs}$ from patient-derived PBMCs measured by LC-MS. Relative abundance is calculated as percentage relative to healthy $T_{regs}$ from healthy blood donors. Patients and controls were age- and sex-matched. n=3 patients/controls; Two-way ANOVA with Holm-Sidak multiple-testing correction *$p<0.05$, **$p<0.01$, ***$p<0.001$. Mean ± SD.

**Extended Data Figure 3**: Blood $T_{regs}$ from patients suffering from chronic skin disease express *SAT1* and have reduced polyamine content.

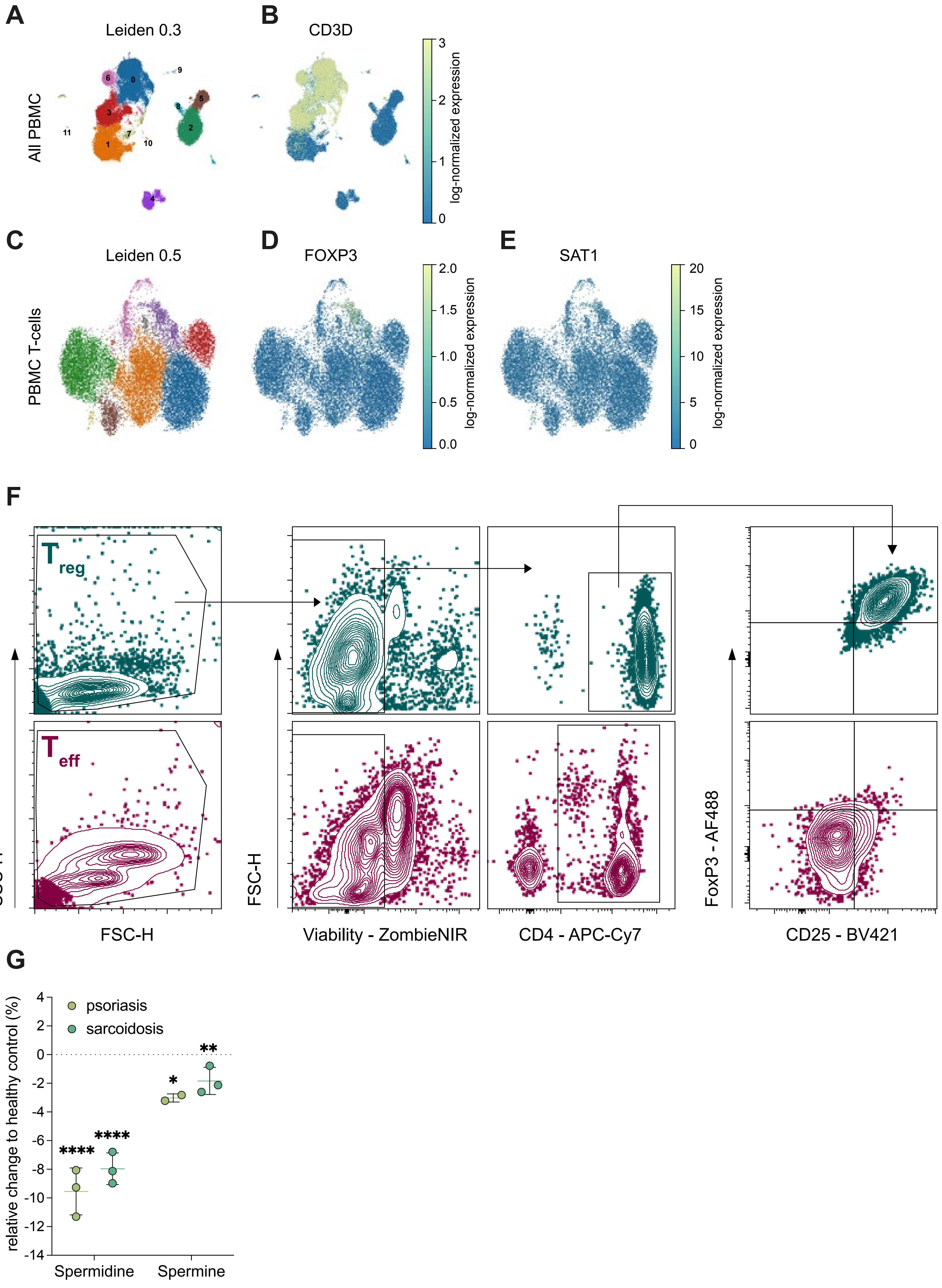

**Extended Data Figure 3: Blood $T_{regs}$ from patient suffering from chronic skin disease express *SAT1* and have a reduced polyamine content.**

**A)** Leiden clustering of PBMCs from healthy donors (n=3) and patients (n=4 psoriasis; n=14 sarcoidosis) with inflammatory skin disease. **B)** UMAP of PBMCs showing expression of CD3D in healthy and patient PBMCs. **C)** Leiden clustering of T cells from healthy and patients with inflammatory skin disease. **D)** and **E)** UMAP of healthy and patient PBMC-derived T cells showing expression of (D) FoxP3 and (E) SAT1. **F)** Flow cytometry analysis of $T_{regs}$ (top) and $T_{eff}$ (bottom) after magnetic isolation based on CD127 and CD25. **G)** Relative abundance of intracellular spermidine and spermine in $T_{regs}$ from patient-derived PBMCs measured by LC-MS. Relative abundance is calculated as percentage relative to healthy $T_{regs}$ from healthy blood donors. Patients and controls were age- and sex-matched. n=3 patients/controls; Two-way ANOVA with Holm-Sidak multiple-testing correction *p<0.05, **p<0.01, ***p<0.001. Mean ± SD.

**Extended Data Figure 4**: $T_{regs}$ in a $T_H2/T_H17$ inflammatory environment have increased *SAT1* expression.

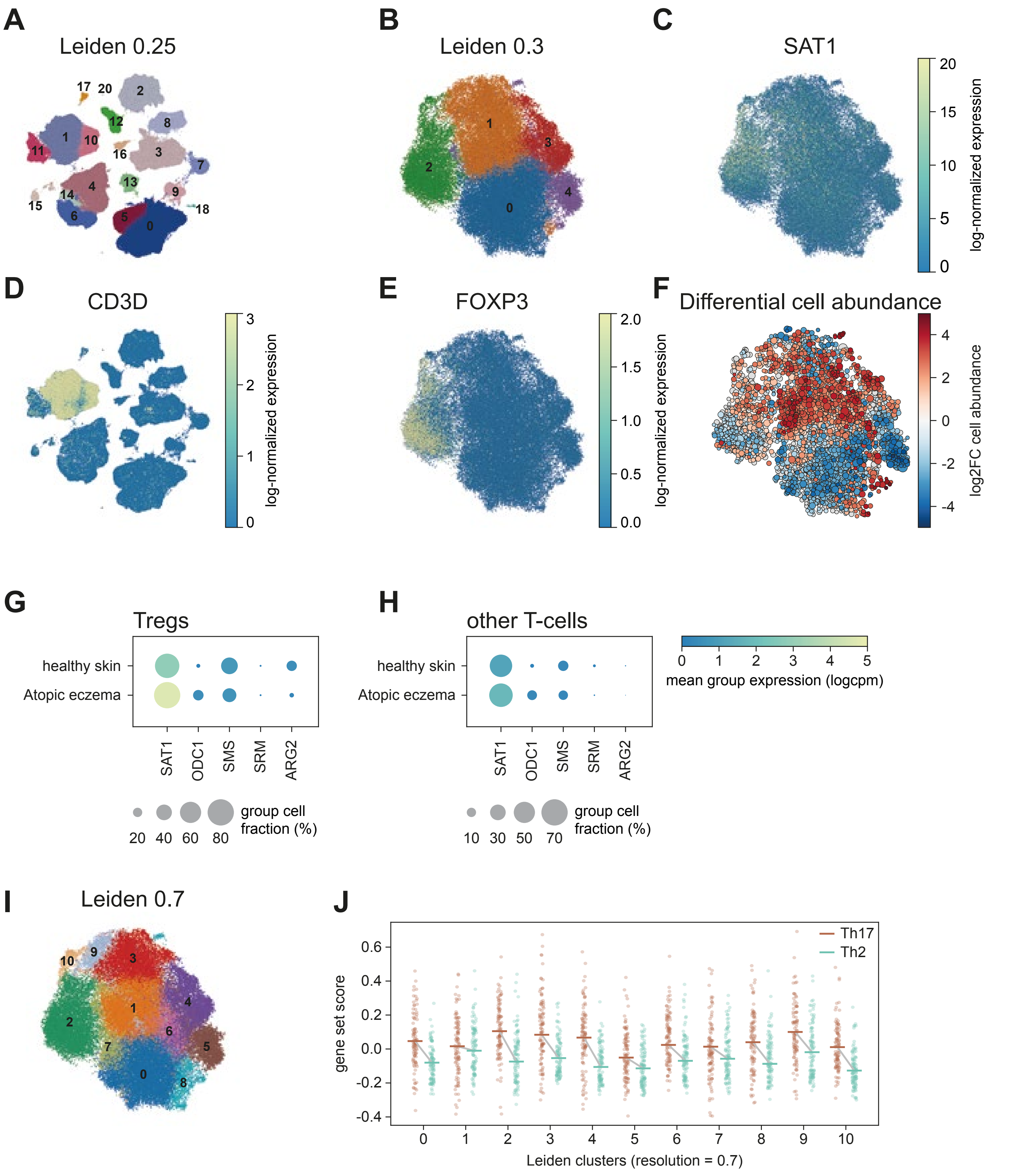

**Extended Data Figure 4: $T_{regs}$ in a $T_H2/T_H17$ inflammatory environment have increased *SAT1* expression.**

**A)** and **B)** Leiden clustering of (A) skin cells and (B) skin T cells from healthy skin and patients with atopic dermatitis. **C)** and **E)** UMAP projection of skin T cells from healthy skin and patients with atopic dermatitis showing (C) SAT1 and (E) FoxP3 expression. **D)** UMAP projection showing CD3D expression in skin from atopic dermatitis and healthy skin. **F)** Differential abundance testing of skin T cells in atopic dermatitis vs. healthy skin. **G)** and **H)** Dotpot of expression of genes involved in polyamine metabolism of (G) $T_{regs}$ and (H) non-$T_{regs}$ in the skin of patients with atopic dermatitis. **I)** Same as (B) but with resolution of 0.7 **J)** $T_H17$ and $T_H2$ scores of clusters in (I) computed as the ratio of average expression of target genes and a random background (see Methods). Line is Median score over all cells in cluster. Dots are a random sample of 100 cells per cluster.

**Extended Data Figure 5**: Skin T cells expand stably over two weeks and are permissive to transduction with nucleofection.

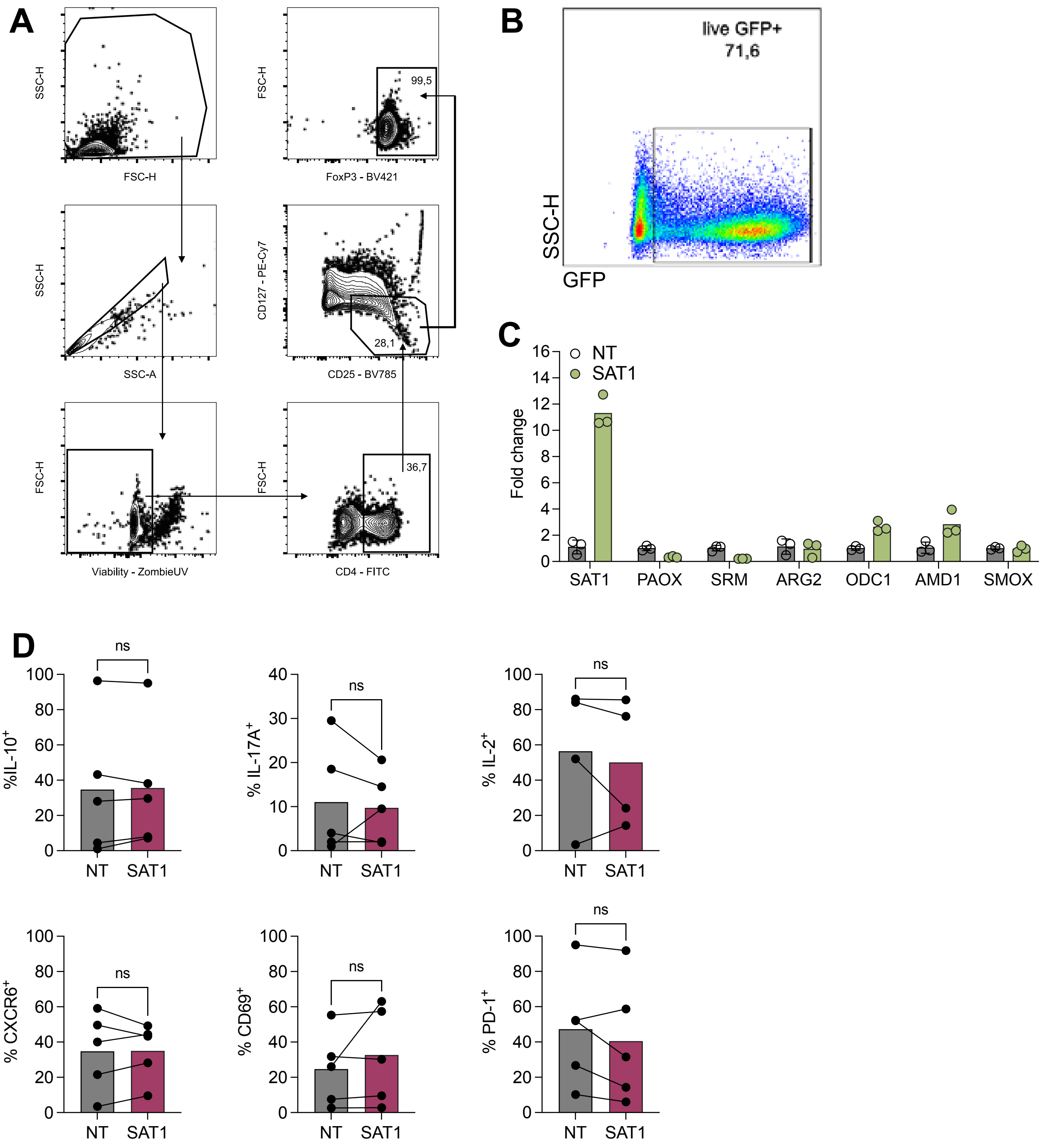

**Extended Data Figure 5: Skin T cells expand stably over two weeks and are permissive to transduction with nucleofection.**

**A)** Representative gating strategy of skin-expanded T cells showing feasibility to isolate skin $T_{regs}$ after explant based on CD127 and CD25. **B)** Transduction efficiency based on GFP expression of skin T cells 24h after nucleofection. Gated on live cells. **C)** Fold change of expression of genes involved in polyamine metabolism in $T_{regs}$ transduced with non-targeting (NT) or SAT1 guide 48h after nucleofection. Mean ± SD. **D)** Quantification of flow cytometry data showing percentage of IL-10$^{+}$, IL-17A$^{+}$, IL-2$^{+}$, CXCR6$^{+}$, CD69$^{+}$, and PD-1$^{+}$ cells in GFP$^{+}$ CD4$^{+}$ $T_{eff}$ cells in non-targeting vs. SAT1 transduced cells. n=5 healthy skin donors, Two-tailed Paired t-test, ns p>0.05.

**Extended Data Figure 6**: $T_H$17 stimulated keratinocytes do not affect CD4⁺ $T_{eff}$ cell expression of *SAT1* and associated markers

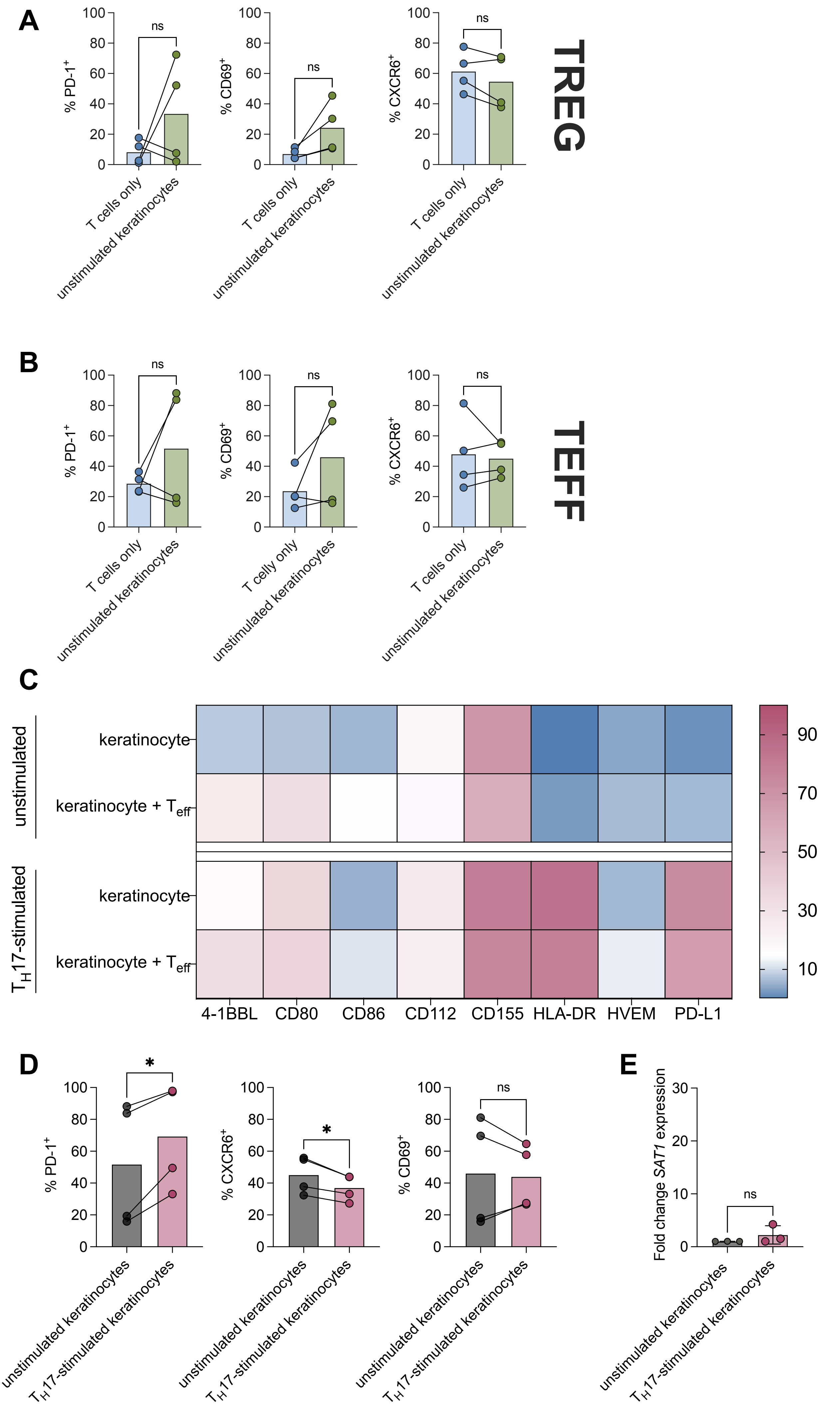

**Extended Data Figure 6: $T_H17$ stimulated keratinocytes do not affect $CD4^+$ $T_{eff}$ cell expression of *SAT1* and associated markers.**

**A)** and **B)** Quantification of flow cytometry data showing percentage of PD-$1^+$, $CD69^+$, and $CXCR6^+$ blood-derived **(A)** $T_{regs}$ and **(B)** $CD4^+$ $T_{eff}$ after 5 days of T cell-only culture or co-culture with unstimulated keratinocytes. **C)** Heatmap of flow cytometry data showing expression of co-receptors by keratinocytes after $T_H17$ stimulation and co-culture with healthy blood-derived $CD4^+$ $T_{eff}$. Scale represents percent of expressing cells. **D)** Quantification of flow cytometry data showing percentage of PD-$1^+$, $CD69^+$, and $CXCR6^+$ blood-derived $CD4^+$ $T_{eff}$ after 5 days of co-culture with $T_H17$ or unstimulated keratinocytes. **E)** Fold change of *SAT1* expression in healthy, blood-derived $CD4^+$ $T_{eff}$ co-cultured with $T_H17$ or unstimulated keratinocytes for 5 days measured by qPCR. Fold change calculated relative to unstimulated control.

**A)-D)** n=5 healthy blood donors, Two-tailed paired t-test, ns $p>0.05$. **E)** n=3 healthy blood donors, Two-tailed paired t-test, ns $p>0.05$. Mean ± SD.

**Extended Data Figure 7**: Co-culture with $T_H2$-stimulated keratinocytes does not induce a SAT1$^{high}$ phenotype in blood-derived $T_{regs}$.

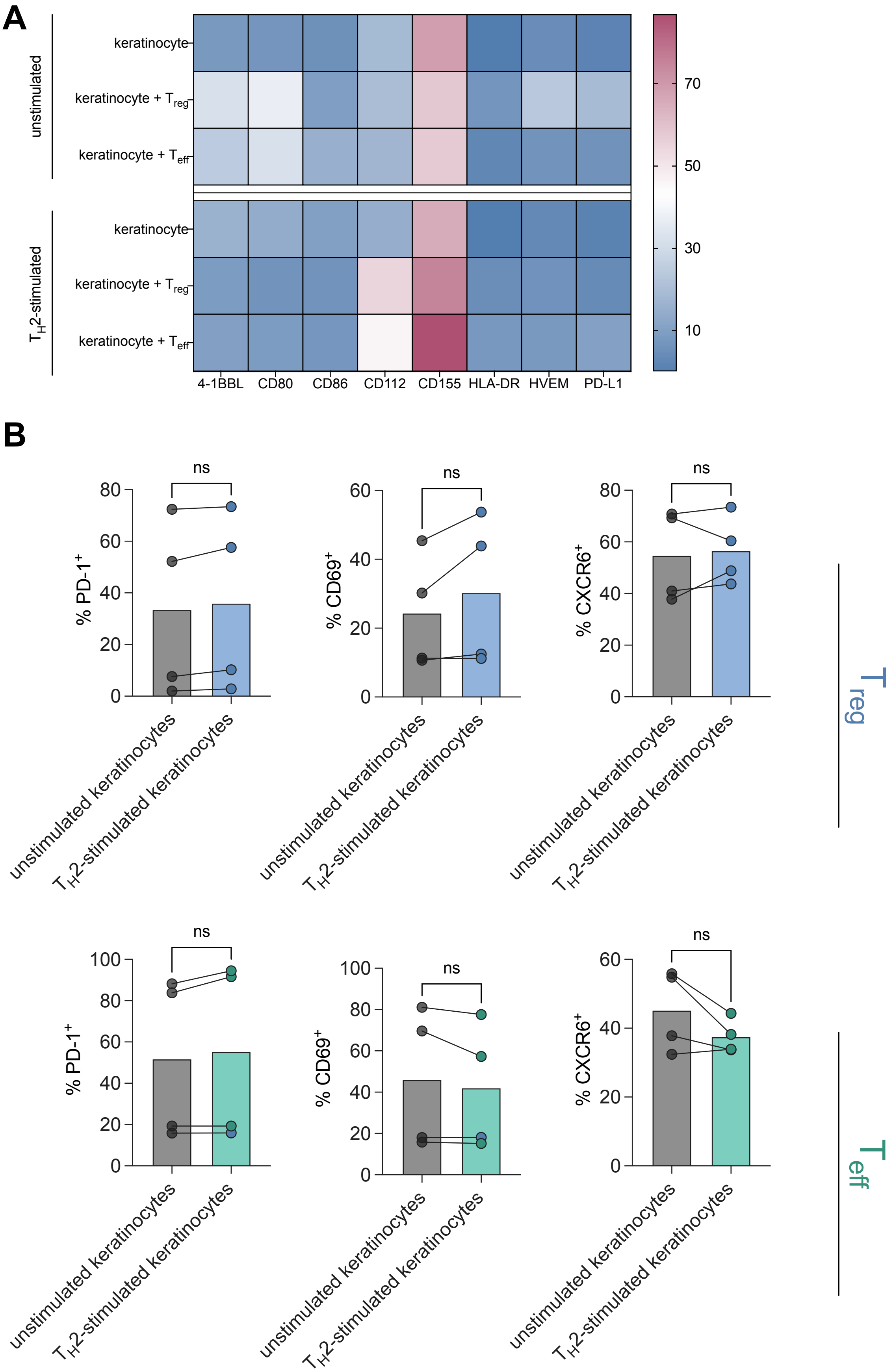

**Extended Data Figure 7: Co-culture with $T_H2$ stimulated keratinocytes does not induce a SAT1$^{high}$ phenotype in blood-derived T cells.**

**A)** Heatmap of flow cytometry data showing expression of co-receptors by keratinocytes after $T_H2$ stimulation and co-culture with healthy blood-derived $T_{regs}$ and CD4$^+$ $T_{eff}$. Scale represents percent of expressing cells. **B)** Quantification of flow cytometry data showing percentage of PD-1$^+$, CD69$^+$, and CXCR6$^+$ blood-derived $T_{regs}$ and CD4$^+$ $T_{eff}$ after 5 days of co-culture with $T_H2$ or unstimulated keratinocytes. n=4 healthy blood donors, Two-tailed paired t-test, ns $p>0.05$.

**Extended Data Figure 8**: High concentrations of diminazene aceturate affect viability and co-receptor expression in keratinocytes.

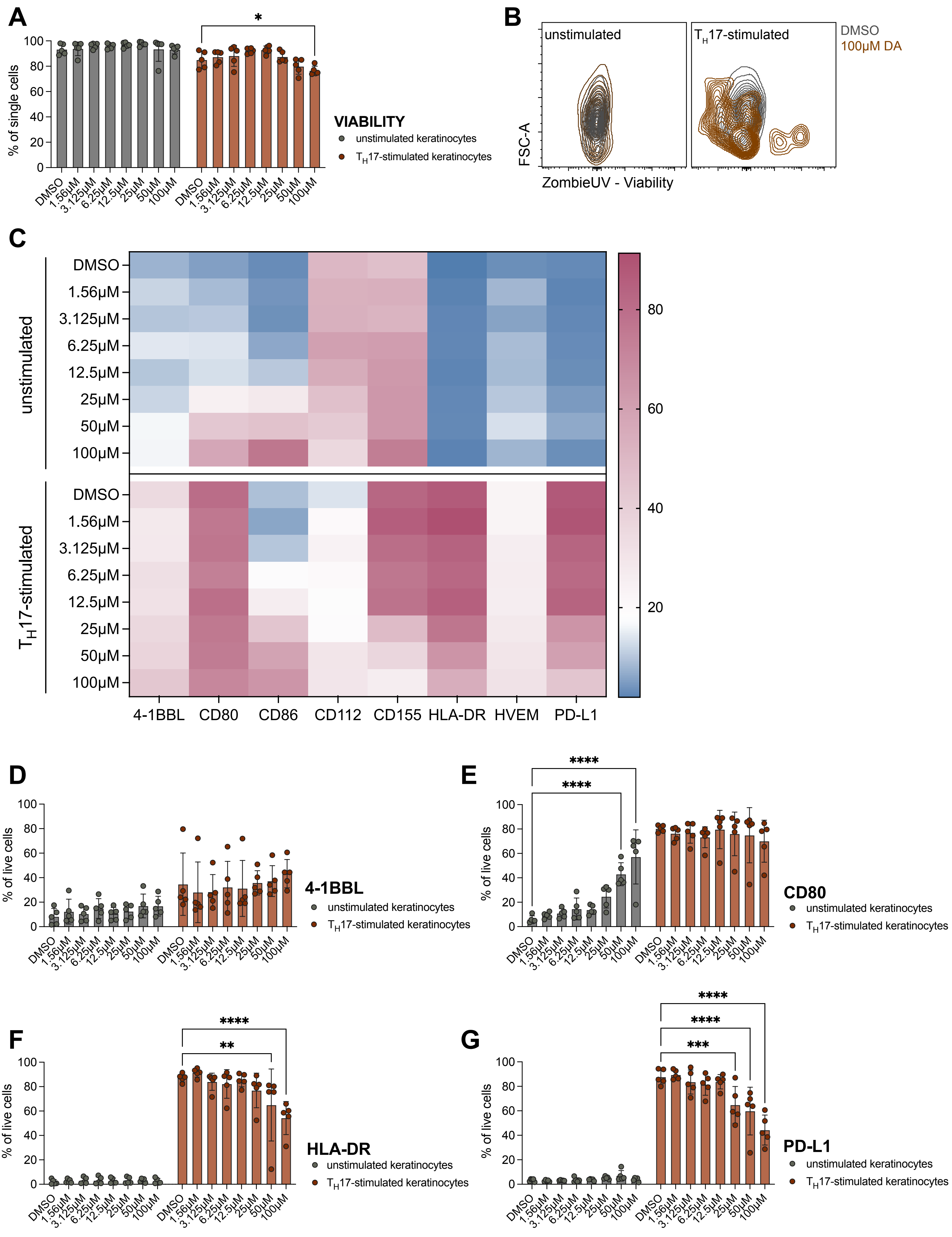

**Extended Data Figure 8: High concentrations of diminazene aceturate affect viability and co-receptor expression in keratinocytes.**

**A)** Quantification of flow cytometry data showing viability of keratinocytes after overnight stimulation with $T_H17$ cytokines or unstimulated control +/- increasing concentration of diminazene aceturate. **B)** Representative flow cytometry density plot showing viability of keratinocytes after $T_H17$ stimulation or control media +/- 100µM diminazene aceturate. **C)** Heatmap of flow cytometry data showing expression of co-receptors by keratinocytes after $T_H17$ stimulation or control media +/- increasing concentration of diminazene aceturate. Scale represents percent of expressing cells. **D)-G)** Quantification of flow cytometry data showing expression of (D) 4-1BBL, (E) CD80, (F) HLA-DR, and (G) PD-L1 by keratinocytes after overnight stimulation with $T_H17$ cytokines or unstimulated control +/- increasing concentration of diminazene aceturate. n=3 separate experiments with technical duplicates, Two-way ANOV Two-way ANOVA with Holm-Sidak multiple-testing correction, *p<0.05, **p<0.01, ***p<0.001, ****p<0.0001. Mean ± SD.

**Extended Data Figure 9**: Diminazene aceturate treatment of DKO$^{K15}$ mice does not affect CD4$^{+}$ T$_{eff}$ cells

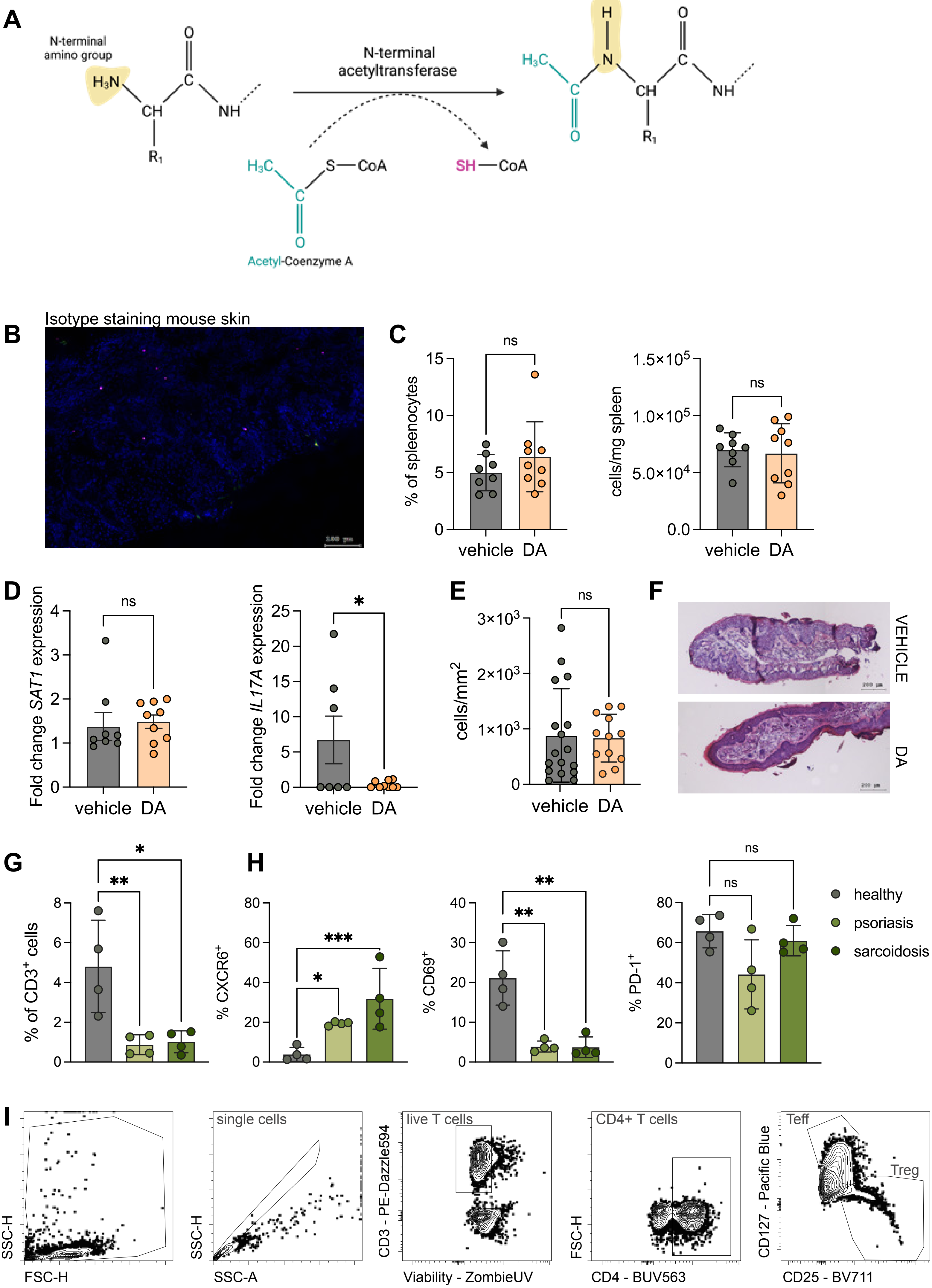

**Extended Data Figure 9: Diminazene aceturate treatment of** DKO$^{*K15}$ **mice does not affect CD4$^{+}$ T$_{eff}$ T cells.**

**A)** Biochemical principle of detection of SSAT enzyme activity in tissue sections. **B)** Isotype staining for $T_{regs}$ in mouse skin. Scale bar = 100µm. **C)** Percentage of $T_{eff}$, and absolute numbers of $T_{eff}$ in the spleen of mice after 10 days with and without treatment. **D)** Fold change of *SAT1* (left) and *IL17A* (right) expression in treated and untreated spleen-derived $T_{eff}$ measured by qPCR. Fold change calculated relative to untreated control. **E)** Quantification of immunofluorescence staining for $T_{eff}$ ($CD3^{+}CD4^{+}CD25^{-}$) in skin of DKO$^{*K15}$ mice as absolute numbers/mm$^2$. Each data point represents one skin section. **F)** Representative H&E staining of mouse ears after 10 days with or without treatment. Scale bar = 200µm. **G)** Quantification of flow cytometry data of patient and healthy PBMCs showing percentage of $T_{regs}$ in the blood. **H)** Quantification of flow cytometry data of patient and healthy PBMCs showing percentage of CXCR6$^{+}$, CD69$^{+}$, and PD-1$^{+}$ $T_{regs}$ in the blood. **I)** Gating strategy for PBMC $T_{regs}$ using CD127 and CD25 expression.

**C)** and **D)** n=8-9 mice per group from 3 different experiments, two-tailed unpaired t-test, *$p<0.05$, **$p<0.01$, ***$p<0.001$. Mean ± SD. **E)** Data from 4 mice per group with 2-3 sections stained per mouse, two-tailed unpaired t-test, *$p<0.05$, **$p<0.01$, ***$p<0.001$. Mean ± SD. **G)** n=4 patients or healthy donors, Two-way ANOVA with Holm-Sidak correction for multiple testing, *$p<0.05$, **$p<0.01$, ***$p<0.001$. Mean ± SD.

| Patient # | YOB | Sex | Time of diagnosis | Diagnosis | Other diagnosis/ Comorbidities | Immunotherapy at time of sampling | Topical therapy at time of sampling |
|---|---|---|---|---|---|---|---|
| HC2 | 1969 | F | NA | Healthy control | None | NA | NA |
| HC2 | 1968 | F | NA | Healthy control | None | NA | NA |
| HC3 | 1980 | M | NA | Healthy control | None | NA | NA |
| SrcX1 | 1974 | F | 2013 | Pulmunar & cutaneous sarcoidosis | None | No | Cortison, Furon, Concor |
| SrcX2 | 1959 | F | 2020 | Cutaneous sarcoidosis | Dyspnoe | No | Clobetasolpropionat cream |
| SrcX3 | 1968 | M | 2019 | Cutaneous sarcoidosis | None | No | Clobetasolpropionat cream |
| Pso1 | 1991 | M | 2012 | Psoriasis vulgaris (PASI > 10) | None | No | NB-UVB |
| Pso2 | 1963 | M | 2017 | Psoriasis vulgaris (PASI > 10) | Smoking | No | No |
| Pso3 | 1991 | M | Unknown | Psoriasis vulgaris (PASI 15.4) | None | No | No |

**Supplementary Table 1: Metadata of patient samples used for metabolomics.**

| **Type** | **Gene** |
|---|---|
| Treg_Trm | IL2RA |
| Treg_Trm | IL7R |
| Treg_Trm | CD69 |
| Treg_Trm | CXCR6 |
| Treg_Trm | CXCR4 |
| Treg_Trm | LGALS3 |
| Treg_Trm | RUNX3 |
| Treg_Trm | RORA |
| Treg_Trm | CCR6 |
| Treg_Trm | CD7 |
| Treg_Trm | MAL |
| costimulation_TCR signaling | TNFRSF4 |
| costimulation_TCR signaling | TNFRSF9 |
| costimulation_TCR signaling | CTLA4 |
| costimulation_TCR signaling | TIGIT |
| costimulation_TCR signaling | ICOS |
| costimulation_TCR signaling | CD28 |
| costimulation_TCR signaling | KLRB1 |
| costimulation_TCR signaling | LCK |
| costimulation_TCR signaling | CD83 |
| costimulation_TCR signaling | SLA |
| costimulation_TCR signaling | TRBC2 |
| costimulation_TCR signaling | CD74 |
| costimulation_TCR signaling | CD70 |
| costimulation_TCR signaling | LY6E |
| cell_death | FAS |
| cell_death | BCL2 |
| cell_death | BAX |
| cell_death | BID |
| cell_death | GPX4 |
| cell_death | TNFAIP3 |
| cell_death | BIRC3 |
| cell_death | HSPA1A |
| cell_death | HSPB1 |
| cytokine_signaling | STAT3 |
| cytokine_signaling | STAT5A |
| cytokine_signaling | STAT5B |
| cytokine_signaling | JAK1 |
| cytokine_signaling | IL1R2 |
| cytokine_signaling | IL21R |
| cytokine_signaling | SOCS1 |
| translation | EIF1 |

**Supplementary Table 2: Selected differentially expressed genes in SAT1$^{high}$ vs. SAT1$^{low}$ cells.**

GRAPHICAL ABSTRACT

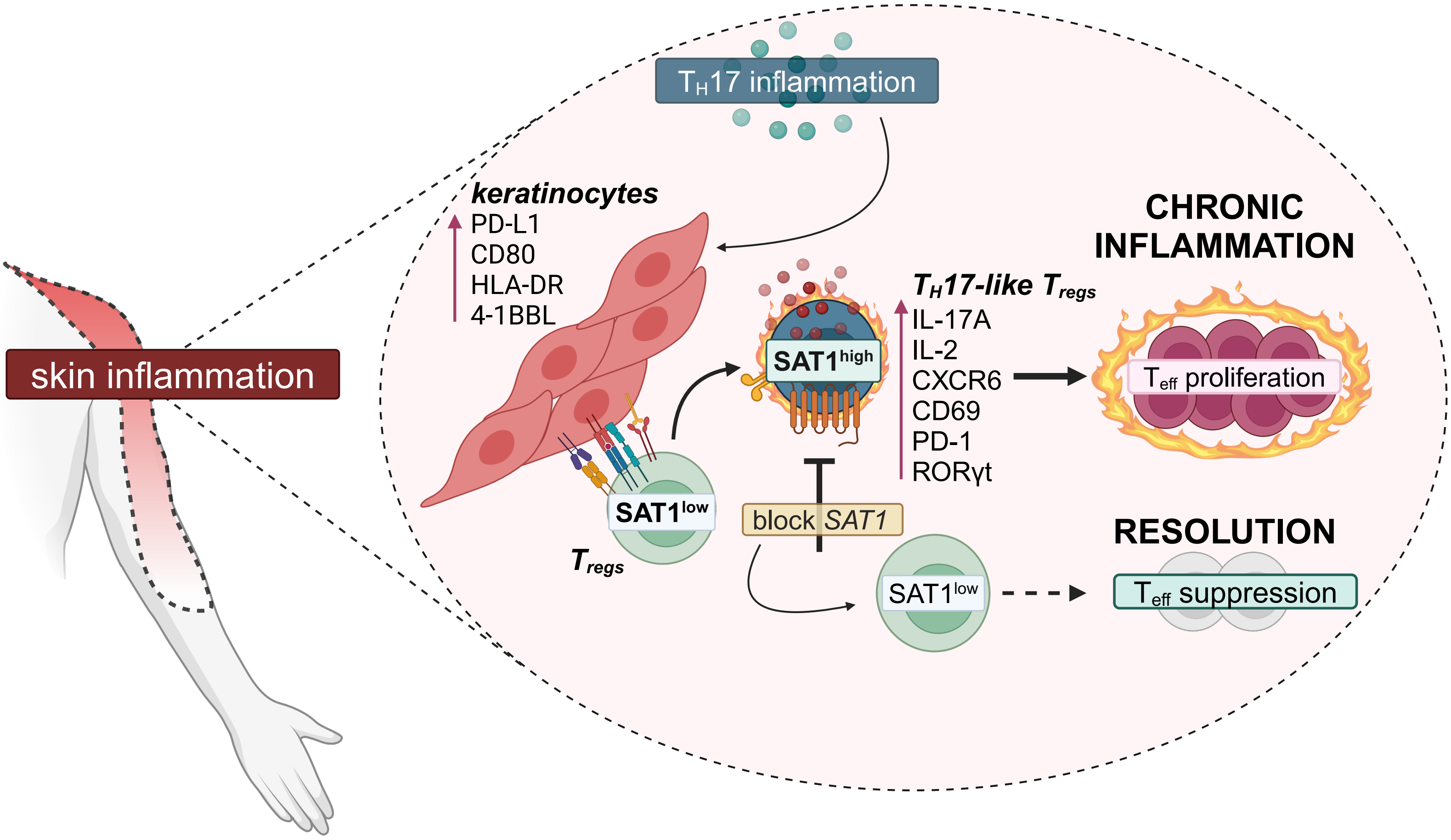

**Graphical abstract**

Created using BioRender.
